# Pollen sterols are highly diverse but phylogenetically conserved

**DOI:** 10.1101/2025.04.21.649781

**Authors:** Ellen C. Baker, Ellen Lamborn, Julia Quiñonez, Stefania Karlsdottir, Abigail Sheppard, Elynor Moore, William Kunin, Geraldine A. Wright, Philip C. Stevenson

## Abstract

Sterols regulate cell membrane fluidity and are precursors for hormone and secondary metabolite production in plants, but plant sterols also have a critical role as nutrients in herbivores. Here we describe the distribution of 78 different sterols from pollen of 295 UK wildflower taxa and use this data to develop an evolutionary rationale for the diversity of sterols in pollen compared to vegetative tissues. The sterolome was a function of plant lineage and conserved in groups as high as subfamily. Insect herbivores are auxotrophic for sterols, and notably bees can not modify them therefore rely on dietary sources, primarily pollen, to meet their metabolic needs. Most pollen in the present study contained high proportions of Δ5 sterols including β-sitosterol, 24-methylenecholesterol and isofucosterol, which are important sterols for bees. The sterols recorded in honey bees occurred in the pollen of only 68% of plant taxa, however, none matched these proportions exactly suggesting they must forage pollen from multiple plant taxa to satisfy their sterol requirements. We conclude that there is evidence for pollen sterol composition being the result of diverse driving forces including plant lineage and pollinator nutritional requirements.

## Introduction

Sterols are lipids which serve as essential cell membrane components. In vertebrates, cholesterol is dominant whereas fungi primarily use ergosterol. Plants, however, produce a wide range of sterols with over 250 reported (Valitova, Sulkarnayeva and Minibayeva, 2016; Nes, 2011). While the most common sterols in vegetative tissues are β-sitosterol and stigmasterol (Behmer and Nes, 2003; Dufourc, 2008) pollen contains a more diverse range of sterols (Zu *et al*., 2021). Some pollen, for example, has no β-sitosterol or stigmasterol but instead contains other sterols such as 24-methylenecholesterol (Vanderplanck *et al*., 2014; Villette *et al*., 2015; Moerman *et al*., 2017; Vanderplanck, Zerck, *et al*., 2020; Zu *et al*., 2021; Furse, Martel, *et al*., 2023). The extent to which the sterolome of angiosperm pollen varies and the adaptive function for this variation remains largely unknown.

Pollen sterols facilitate fertilization of the ovule by providing substrates for pollen tube elongation (Rotsch *et al*., 2017; Stroppa *et al*., 2023). Sterols are especially important for rapid cell elongation and cell polarisation during pollen tube growth (Rotsch *et al*., 2017). However, the sterols used for pollen tubes are not those occurring in mature pollen grains (Rotsch *et al*., 2017) but are created *de novo* during pollen tube growth (Rotsch *et al*., 2017; Stroppa *et al*., 2023). Since sterols are also metabolically expensive (Nes, 2011) and in the absence of an established explanation of their function, it is surprising that they accumulate in pollen with such diversity.

A factor that could influence sterol diversity in pollen is the nutritional needs of pollinators such as bees (Rotsch *et al*., 2017). Like other insects bees cannot synthesize sterols *de novo* but instead acquire them along with protein, fats and micronutrients from their diets (Wright, Nicolson and Shafir, 2018). However, unlike other phytophagous insects, bees cannot convert dietary sterols to cholesterol (Behmer and Nes, 2003). Instead, they incorporate specific pollen sterols directly into their tissues without modifying them (Svoboda *et al*., 1986). The sterol profile of the few bees that have been studied (honeybees, leaf-cutting bees) is mainly composed of 24-methylenecholesterol, isofucosterol, β-sitosterol, campesterol, desmosterol and cholesterol (see Svoboda and Feldlaufer, 1991, Moore et al., 2025). The use of pollen-specific sterols in bee tissues may be an adaptation to pollinivory (Svoboda, Feldlaufer and Weirich, 1994) but would be an evolutionary risk if pollen that bees collected did not contain all the required sterols.

Anthropogenic change to flowering landscapes throughout the world has significantly altered floral resource availability for pollinators (Howard *et al*., 2003; Vanbergen and Insect Pollinators Initiative, 2013; Baude *et al*., 2016), increasing the likelihood of nutritional deficiencies. An understanding of pollen sterol variation could inform predictions about nutritional stress in bees. To measure sterol production in pollen, we measured the diversity and abundance of sterols in 295 UK wildflower taxa using a sterol analysis. We found that sterol profiles were remarkably stable within taxonomic groups, including subfamilies of Asteraceae and Rosaceae. Sterolomes did not vary at different sites and a significant phylogenetic signal for almost half of the sterols was recorded. However, none of the pollen sterolomes matched those reported from honeybees.

## Methods

### Species selection

In total, 700 pollen samples were collected for sterol analysis representing 295 taxonomic units, 275 identified to species level, and 125 of which were represented by a single sample. A full summary of samples collected, dates and location are provided in Supplementary Table 11. Samples included pollen from 172 species selected from the species list compiled by (Baude et al., 2016) which targeted plants from the Countryside Survey in Great Britain that were rewarding to pollinators and considered sufficiently common to contribute to landscape level floral resources. During collection visits for the above target species, 88 plants which produced easily extractable pollen or would contribute to wider taxonomic coverage in the dataset were also opportunistically collected. Asteraceae flowers produce copious and easily collectable pollen and previous work has suggested nutritional deficiencies within Asteraceae pollen (Vanderplanck, Gilles, *et al*., 2020). Therefore, Asteraceae flowers were also targeted opportunistically adding a further 18 species while 17 more species were targeted due to their importance as forage plants for wild bee species in the UK (such as *Bryonia cretica* for *Andrena florea*, *Reseda* spp. for *Hylaeus signatus* and *Anthyllis vulneraria* for bumble bees).

### Pollen collection

Samples were collected from a range of sites in England including nature reserves, botanic gardens, public land and private gardens from spring to early autumn from 2021-2022. In total, samples were collected from 15 counties. Collection months and counties for all samples are shown in Table 11. Flowering heads, or branch tips in the case of woody species, were cut from the plant and placed in sealed sandwich bags. These were then taken back to the laboratory and placed in a beaker of room temperature tap water to dehisce. The duration needed for anthers to dehisce ranged from a few hours to a few days, but most were left for 24 hours. Once pollen had been produced it was removed from the flower using a clean paintbrush and forceps under a magnifying glass. Pollen was collected onto an ethanol-wiped glass microscope slide and visually inspected for debris or insects which were subsequently removed. The pollen sample was then scraped into a clean 1.5ml Eppendorf tube or glass vial using an ethanol-wiped razor blade. The exact method of pollen extraction varied depending on species and floral morphology. For daisy-type Asteraceae (Asteroideae), pollen was brushed from the tops of disk florets in clumps. For thistle-type Asteraceae (Carduoideae), pollen was brushed from anther tube tips and tubes squeezed gently to remove pollen further. For legume-type Fabaceae flowers, wing petals were removed and the tip of the keel lightly squeezed, as for *Lotus corniculatus,* or all petals removed entirely, as for *Trifolium pratense*. Pollen samples were then frozen at −20°C before being moved to −80°C in batches. Freezing pollen has been shown to prevent nutritional degradation (Hagedorn and Burger, 1968; Pernal and Currie, 2000).

To assess inter- and intra-species variation in sterol profile, a subset of species was collected from at least three counties in England. Plant identification was confirmed by photos shared on iNaturalist and with botanists at Royal Botanic Gardens (RBG), Kew. A total of 275 samples were identified to species and subspecies level and 20 identified to genus level.

A minimum of 10 mg fresh weight pollen per sample was required due to the lower end of pollen total sterol content being 0.1%, i.e., 10 µg of sterol in a 10 mg sample, a level within the detectable range of our analysis. At the end of the collection season all samples were transferred to 1.5 ml glass vials and partitioned or combined by species, depending on whether the sample was over or under 10 mg. The final weight of each sample was between 9.5-16.5 mg.

### Sample preparation

Alongside the hand collected pollen samples, additional samples were used in the sample run to monitor performance. Firstly, Quality Assurance (QA) samples contained a solution of reference materials compiled to assign names to signals in our samples. This allowed us to assess the consistency of analysis by the instrument across a large cohort of samples and calculate the coefficient of variance. Reference materials verified by multi-nuclear NMR were 24-methylenecholesterol, 24-methylenecycloartenol, anthelsterol (a sterol found in *C. anthelmintica*, ST(28:3), that is active under 330 nm UV but whose structure has not been determined formally using NMR (Furse, Martel, *et al*., 2023), avenasterol, β-sitosterol, brassicasterol, campesterol, cholesterol, cycloartenol, cycloeucalenol, cyclolaudenol, desmosterol, episterol, ergosterol, isofucosterol, schottenol, sitostanol, spinasterol and stigmasterol. The *m/z* and retention time for each sterol is shown in Supplementary Table 12. Aliquots of 40 µl were used in the run. Secondly, a Quality Control (QC) stock solution was made by homogenizing a set of commercially available honey bee collected pollens: 95 g of *Helianthus annus* (sunflower) pollen, 5 g of *Nymphaea* sp. pollen, 20 g of *Fagopyrum esculentum* (buckwheat) pollen in 120 ml water. For this project three QC concentrations were used: 100%, 50%, 25%, corresponding to 40 µl, 20 µl and 10 µl aliquots of the QC stock solution. These were analysed in the same way as the samples to calculate the correlation between analyte concentration and signal size. Variables whose ratio was <0.75 were discarded. All runs began with a blank containing LCMS quality methanol only. This was designed to enable identification and removal of background noise.

### Sterol extraction

To each vial of hand collected pollen or QC sample a 20 μl aliquot of 0.1 mg/L (5.084 pMoles/sample) *d_7_*-cholesterol as the internal standard (IS) was added with 300 µl methanoic KOH (10 g:90 ml) and the vial vortexed for 20 seconds. Vials were then placed in drilled aluminium heating blocks at 75°C in a water bath for two hours. Saponification releases conjugated sterols (Evtyugin *et al*., 2023), such as sterol esters which are reported to be common in pollen (reviewed in Ferrer *et al*., 2017). The saponified material was then transferred to a corresponding deep 96-well plate (96-well plate, Esslab Plate+™, 2.4 ml/well, glass-coated). Pollen samples were randomised within plates and interspersed randomly with QA, QC and blank (300 µl methanol and 20 µl IS) samples.

Extraction of the sterol fraction was carried out on the plate using a 96-channel pipette (liquid-handling robot, Integra Viaflow 96/384 channel pipette) using DMT (dichloromethane (DCM) (3 parts), methanol (1 part) and triethylammonium chloride (0.0005 parts, i.e. 500 mg/l). To each well 500 µl of DMT (250×2) was added followed by a 500 µl aliquot of deionised water and the mixture agitated using the 96-channel pipette. The plate was centrifuged for 2 min to separate layers (methanol + water, solid, DMT + sterol). A 50 µl sample of extract (DMT & sterol) was transferred to the 384-well plate and left for 30 min for the DMT to evaporate. After repeating this process four times, 150 µl of LCMS grade methanol was added and the plate sealed with foil. A total of three 384-well plates were run immediately in sequence, analysing 700 hand collected pollen samples and 281 analytical samples. One pollen sample was later excluded because it had experienced prolonged exposure to room temperature conditions during collection.

### Sample analysis and data processing

Mass spectra were recorded using a Thermo Scientific Orbitrap Fusion mass spectrometer, which was equipped with an Atmospheric Pressure Chemical Ionization (APCI) probe and integrated with a Thermo Scientific Vanquish liquid chromatography system (Thermo Scientific, UK). Metabolites were separated prior to analysis on a Hypersil GOLD LCMS C18 column (50 × 2.1 mm with particle size 1.9 μm) (Thermo Scientific, UK) which was maintained at 55 °C for analyses. An aliquot of 1 μL was injected, with a mobile phase flow rate of 0.60 mL/min comprising a gradient of 84% methanol at T = 0 to 100% methanol at T = 11 min and held at 100% till T = 16.4 min, and then returned to 84% methanol from 16.5 to 19 minutes. Mass spectrometric data were acquired in positive ionisation mode via APCI, operating at a resolution of 120,000. The spray voltage was set to 2.86 kV, with sheath nitrogen gas flows of 45 (sheath), 5 (auxiliary), and 1 (sweep) arbitrary units. The ion transfer tube and vaporizer temperatures were maintained at 300 °C and 350 °C, respectively. The mass acquisition range was calibrated to *m/z* 197–500, with the lower limit adjusted to detect the fluoranthene cation (*m/z* 202.077), which served as the internal mass calibration reference. Under these conditions, the internal standard used for the samples, *d_7_*-cholesterol, eluted at 5.6 minutes, with the primary ion species identified as [M - H_2_O + H] + at *m/z* 376.395 within a mass variance of less than 1 ppm. Both samples and external standards were analysed in a single sequence. The pollen samples were randomised among themselves, while the external standards were run sequentially, beginning with the least concentrated standard. Instrument output was processed after Furse, Martel, *et al*. (2023). Data files were processed using Analyzer PRO XD (SpectralWorks Ltd, Cheshire, UK). Signals were extracted from 3 to 11 min in a mass range of 360–442 *m/z*, with an area threshold of 500 000, smoothing of 15 arbitrary units, a width of 0.01 min, mass accuracy of 3 d. p. and a retention time window of 0.1 min.

All compound signals were determined by their *m/z* and retention time (level 3 annotation) 78 of which were further distinguished using the format ST (27:1)A where ST represents ‘sterol’, the first number indicates number of carbons, the second number indicates number of double bonds and the letter distinguishes sterols that differed but otherwise had a similar number of carbons and double bonds (level 2 annotation). Compounds were assigned names (a level 1 annotation) where reference materials that had been verified using multi-nuclear NMR were available as reference samples. These included 24-methylenecholesterol, 24-methylenecycloartenol, agnosterol, anthelsterol, avenasterol, β-sitosterol, brassicasterol, campesterol, cholesterol, cycloartenol, cycloeucalenol, cyclolaudenol, desmosterol, episterol, ergosterol, isofucosterol, lophenol, schottenol, sitostanol, spinasterol, and stigmasterol. Anthelsterol is a trivial name for a sterol that has not been determined using NMR (Furse et al., 2023).

Instrument output reported mass to charge ratio (*m/z*), signal size and retention time. Signal size output was first converted by dividing the signal area for each metabolite by the signal of internal standard (*d_7_*-cholesterol). These relative abundance values were used for the majority of the data analysis. To generate a value of total sterol production in sterol/fresh weight of pollen in mg/kg the following formula was used:

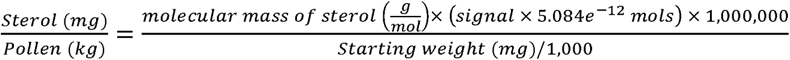

### Data analysis

All data analysis was done in R version 4.4.0 (2024-04-24 ucrt) (R Core Team, 2024) and R studio (2024.9.1.394) (Posit team, 2024). A total of 78 sterols and related compounds were identified in the sample set, and a total of 295 taxonomic units were analysed including those identified to genus, species and subspecies level. Summary statistics were calculated using means of these 295 units with some then removed for species level and phylogenetic analyses as detailed below.

Outliers were identified using disana() from the LABDSV package v.2.1-0 (Roberts, 2023) on a Bray-Curtis dissimilarity matrix of proportion data from all samples. Any data point with a minimum dissimilarity >0.5 compared to all other points in the data was identified as an outlier, as per author guidance, and so removed. This led to the removal of one *Campanula trachelium* sample. Simpsons diversity index was calculated using diversity(index=“simpson”) from the VEGAN package v.2.6-6.1 (Oksanen *et al*., 2024) for all 295 taxonomic units to better understand differences in sterol diversity between pollens.

NMDS were plotted using metaMDS(trymax=200, k=2, autotransform=FALSE). The lowest number of axes was selected where stress was ≤0.2 (established cut off, (Bakker, 2024)). Stress values reflect how well the ordination aligns with the original data. Proportion data (0-1) was used for analysis, where zero denotes a sterol is absent from a sample, and one denotes it is completely dominant. Proportions were arcsine square root transformed and converted into a Bray-Curtis dissimilarity matrix using vegdist(method=”bray”). Where NMDS stress value was <0.2 a 2D biplot as made using GGPLOT2 v.3.5.1 (Wickham, 2016) and GGREPEL v.0.9.5. (Slowikowski, 2024) to show significant (*p<*0.010) and strong (>0.7 absolute value) associations. PERMANOVA was calculated using adonis2() to test for significant differences between groups. Betadisper() and anova() were used to confirm non-significant difference in dispersion. Adonis2() was also used to test for pairwise comparisons between groups following Bakker (2024), using a Benjamini-Hochberg correction which corrects for multiple testing to reduce the chances of false positives. All functions were from VEGAN (Oksanen *et al*., 2024) except anova() from the STATS package (R Core Team, 2024). The STATS package from R (R Core Team, 2024) was used to calculate summary statistics. Indicator Species Analysis (ISA) was carried out on arcsine square root transformed data to accompany NMDS plots using multipatt() from the INDICSPECIES package v.1.7.14. (De Cáceres and Legendre, 2009) to determine significant associations between groups and individual sterols. ISA was done at the level of individual groups only and not for combinations of groups.

### Phylogeny construction and phylogenetic signal

A phylogenetic tree of the study species was constructed based on the date calibrated and rooted tree from Zanne *et al*. (2014). The tree was pruned for our study species using keep.tip() from the R package APE v.5.8.(Paradis and Schliep, 2019). Any synonyms present in the tree were manually renamed to match the POW taxonomy used in the project (19 species). In the case of species which did not have a corresponding match on the Zanne phylogeny the following approach was taken: Where a subspecies level match was not available, a species level branch was renamed instead (2 species). Where a hybrid species was not available, a matching parent not already in the dataset was renamed (5 species). Where a species match was not available, another species from the same genus was renamed instead (7 species), but only where there was not already a species correctly matched within that genus. Out of 295 taxonomic units, 34 could not be suitably placed on the tree and were removed from phylogenetic analysis.

Families were checked for monophyly using AssessMonophyly() from the R package MONOPHY v.1.3.1.(Schwery and O’Meara, 2016). Phylogeny was plotted with ggtree() from GGTREE v.3.12.0.(Yu *et al*., 2017, 2018; Yu, 2020; Xu *et al*., 2022). Heatmap and bar graph associated with phylogeny were plotted using ggplot() from TIDYVERSE v.2.0.0.(Wickham *et al*., 2019). The composite graph was then plotted using APLOT v.0.2.2. (Yu, 2023).

Of the 78 sterol compounds detected, the 69 that had a maximum proportion >1% were analysed for phylogenetic signal (Zu *et al*., 2021). To determine if there was a phylogenetic signal in sterol production across the data, whereby closer related species would share more similar profiles than expected by chance, Pagel’s λ and Blomberg’s K were calculated. Both metrics use a Brownian model of trait evolution and in both cases a value of 0 indicates the trait has evolved independently of phylogeny, therefore closely related species are not more similar to each other than distantly related ones. Low phylogenetic signal can mean high rates of evolution are causing big differences between closely related species or perhaps convergent evolution is causing distant relatives to be more similar than close ones. High phylogenetic signal (K, λ=1) can be caused by low rates of evolution or stabilising selection. Interpretation of high and low phylogenetic signal varies between studies (Kamilar and Cooper, 2013). Phylo4d objects were generated from species tree and proportion data using phylo4d() from PHYLOBASE v.0.8.12.(R Hackathon et al., 2024) and phyloSignal() from the R package PHYLOSIGNAL v.1.3.1.(Keck *et al*., 2016) used to calculate Pagel’s λ and Blomberg’s K. The number of iterations was set to 999. Quantitative values (mg/kg) were used for total sterol content and proportions (0-1) for all other sterol traits. To identify trends in different types of sterols, proportion data was also grouped according to carbon count and position of the B-ring double bond. Ergosterol was classed as NA for double bond position as it contains double bonds at positions 5 and 7 and sterols with a cyclopropane ring were classed as CPR. Un-named sterols were all classed as ΔNA, full details are available in Table 13.

### Family, genus and species level sterol diversity

Prior to analysis of inter-species differences, 17 samples identified to only genus level where there was already a species level representative in the genus were removed.

Species within the family Asteraceae were grouped according to subfamily: Asteroideae (n=21, tubular and ray florets, Cichorioideae (n=17, ray florets only), Carduoideae (n=14, tubular florets only). Subfamily classifications were taken from the Open Tree of Life version 14.7 (OpenTreeOfLife *et al*., 2019) and (Mandel *et al*., 2019). Mean sterol proportions were calculated for each species. An NMDS was plotted containing all Asteraceae species (stress = 0.166) to compare sterol profile between the subfamilies and an ISA carried out to test for associations between subfamilies and specific sterols.

For each species, sterols were then grouped by B-ring double bond saturation (CPR, Δ0, Δ5, Δ7, Δ8 or ΔNA) and their proportions summed (double bond position data is available in Table 13). Ergosterol was classed as NA for double bond position as it contains double bonds at positions 5 and 7. Un-named sterols were all classed as ΔNA. Proportion data (0-1) was arcsine square root transformed and a Kruskal-Wallis rank sum test carried out using kruskal.test(),for each double bond group, followed by a post-hoc Dunn test using dunnTest() from the FSA package 0.9.5. (Ogle *et al*., 2023) to test for differences between Asteraceae subfamilies and the rest of the dataset. Both analyses used a Benjamini-Hochberg correction for multiple testing.

Three families (Asteraceae, Fabaceae and Rosaceae) contained three genera of at least three species each and so were used to analyse intra-family variation in sterolome at the genus level. An NMDS and PERMANOVA were computed separately for the three families using data from individual samples to test for significant differences in sterolome between genera. Species identity was also included in the models.

A total of 48 species were collected from at least three counties in England. As 48 groups would be too many to separate on an NMDS, the species with the highest number of samples were chosen for statistical comparison. Species which were sampled from four different counties and had a least five samples overall were used. An NMDS and PERMANOVA were used to test for significant differences between these most well sampled species (stress value=0.096).

### Ecological associations of sterol profiles

In order to establish how many plants could suitably meet the sterol requirements established for honey bees, data for 18 prepupal honeybee samples (15 prepupae per sample) was used from Svoboda *et al*., (1980) and compared to all hand collected pollen species in this dataset. In order to compare these datasets, only sterols detected in the Svoboda work were used (24-methylenecholesterol, isofucosterol, β-sitosterol, campesterol, cholesterol and desmosterol) and pollen sterol profiles were not recalculated following this removal. An NMDS (k=3) and PERMANOVA were used to show differences in sterol profile composition and variation between the bees and pollens (stress = 0.162).

### Total sterols

For families which contained at least three genera, species data were used to create a box plot showing range in total sterol production (mg/kg) across families.

To compare the effect of collection county on total sterol production, data was subset as follows: Total sterol (mg/kg) was calculated for all samples, data was filtered to only include species which had been collected from at least three counties. Counties were then ranked by the number of these species they included and those with ≥10 kept (Oxfordshire (33), Greater London (25), County Durham (21), North Yorkshire (20), Sussex (10)). This led to a total of 35 species across five counties being included in the analysis. A linear mixed-effects model was created with total sterol production (mg/kg) as the response variable and collection county as a fixed effect. Species identity was included as a random effect. Where a species had been collected more than once from a county the mean value was used for that county. Total sterol data was log_10_ transformed in order to reduce non-normality of the residuals. Model was run using lme() from the NLME package v.3.1.164. (Pinheiro, Bates and R Core Team, 2023). The residuals of the fixed effects showed normal distribution (Shapiro-Wilk test: w=0.986, *p>*0.100) and homogeneity of variance (Levene test: F_4,104_=0.685, *p>*0.500). Random effects also showed normal distribution (Shapiro-Wilk test: w=0.962, *p>*0.100).

## Results

### Quantity of pollen sterol production varies among plant species

We observed interspecies variation in total sterol content of greater than two orders of magnitude. For example, the dandelion, *Taraxacum sect. Taraxacum,* contained the highest amount of sterol (75,423.52 mg/kg sterol/fresh weight pollen). The pollen with the lowest concentration of sterol was observed in the common mallow, *Malva sylvestris,* (729.98 mg/kg, Supplementary – Table 9). Asteraceae accounted for some of the highest and lowest sterol amounts, specifically, Cichorioideae subfamily flowers showed a consistently higher total sterol production than the other Asteraceae flowers (Supplementary - Table 9). Brassicaceae showed the highest sterol content on average, as Asteraceae was still dominated by low sterol producing species (Supplementary - Figure 4).

In general, location of collection did not have a significant impact on total sterol production (mg/kg) (F_4,70_ = 1.254, *p>*0.100). However, variation in total sterol production was still present, potentially caused by abiotic factors, such as temperature, affecting total fat production, or a result of handling and storage procedures working at mg scale. Mean and standard deviation for all species replicated at the county level are reported in Supplementary Table 10.

### Pollen shows a diverse but phylogenetically constrained sterol profile

Within the 295 taxonomic units we sampled (including genera, species and subspecies), we recorded 78 sterols of which 24 were fully assigned using reference compounds (level 3 annotation) and 51 were identified putatively and each determined to be structurally distinct based on chromatography, number of carbons and degree of saturation (noted as ST(carbon number: degree of saturation) from here onwards) (level 2 annotation). Most sterols were present in small proportions even if they dominated in some pollens e.g., desmosterol (median 0.02%, max 83.50%) and cholesterol (median 0.01%, max 61.67%) (Supplementary - Table 1). Thirty-eight sterols were not recorded at higher than 5% in any species. The only sterols with a median of >2.5% were β-sitosterol, isofucosterol and avenasterol (Supplementary - Table 1).

Only four sterols: campesterol, cycloartenol, ST(30:2)B and ST(30:2)C were detected in all taxa and varied in maximum abundance, from 30.23% in campesterol up to 77.92% in cycloartenol, both of which had a median of >1% (Supplementary - Table 2). Cyclolaudenol, ST(30:2)A, ST(30:2)E and ST(31:2)A were in all but one taxonomic unit. Avenasterol, 24-methylenecholesterol and schottenol were in all but two.

Ergosterol, a sterol restricted to fungi (Weete, 1973), was detected in 253 taxonomic units suggesting the near ubiquitous occurrence of fungi in pollen indicating that fungi are inadvertently likely to be a significant part of the diet of pollen feeding animals such as bees. It was not recorded at >1.33% in any species (maximum amount in honesty, *Lunaria annua*). The maximum in an individual sample was recorded in meadow buttercup (*Ranunculus acris,* 1.40%) and had a median proportion across the dataset of 0.01%. Of the 12 samples where ergosterol occurred at >0.75%, nine were *Ranunculus* spp. On average, sampled taxa had 62 sterols in their pollen. The smallest range of sterols (35) was recorded in the pollen of creeping cinquefoil (*Potentilla reptans),* while the largest range was recorded in black alder (*Alnus glutinosa),* which contained 76 sterols. Species also showed a wide variation in sterol diversity as calculated by Simpson’s diversity index. The most diverse species was *Papaver cambricum* (index = 0.913) and the least was *Torminalis glaberrima* (index = 0.169) (*Supplementary* – Table 3, Figure 1).

**Figure 1.**
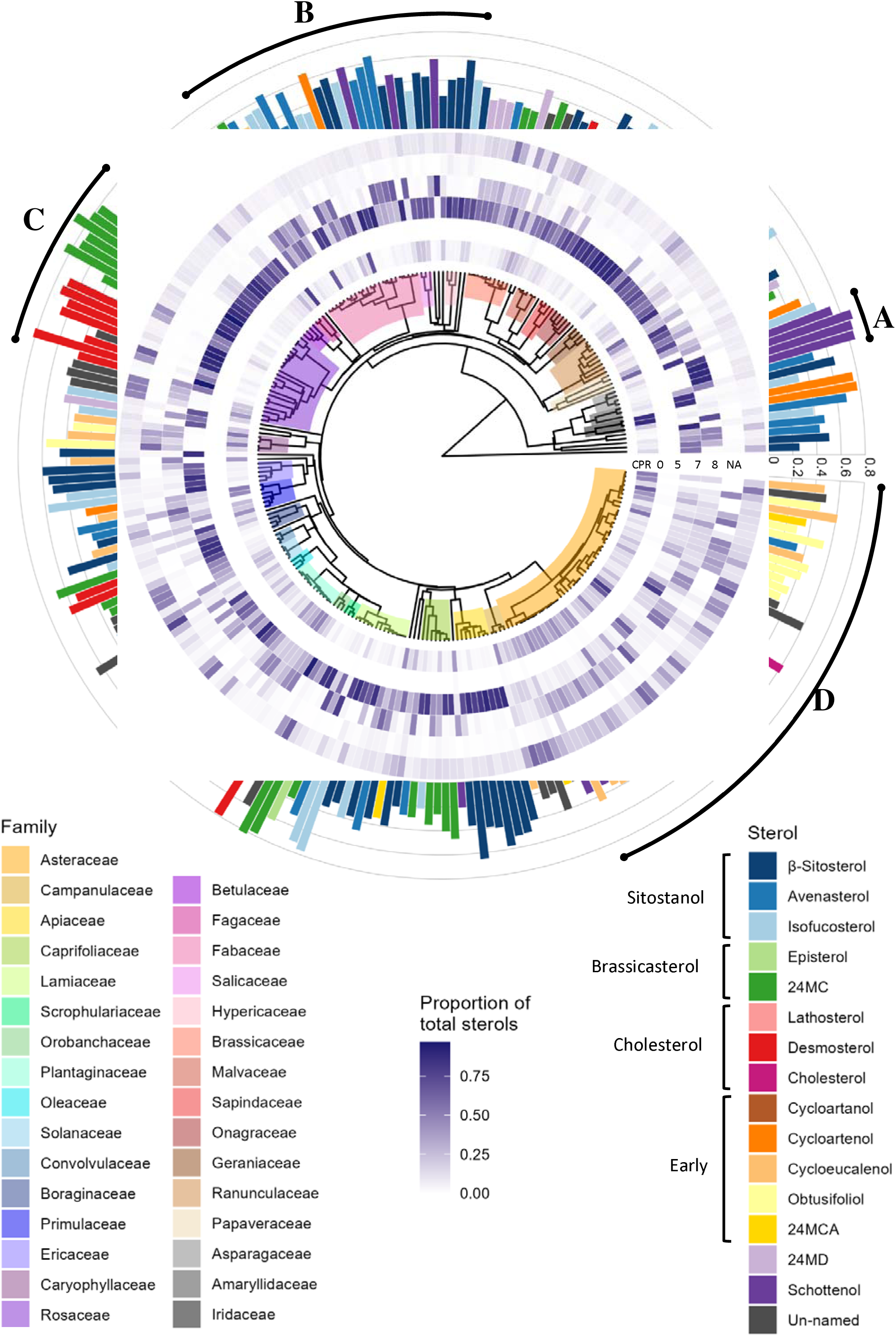
Species phylogeny showing sterol composition across 261 species. Phylogeny is adapted from (Zanne et al., 2014) and all families with three or more species are highlighted, reading from the right in a clockwise direction. The inner rings show a heatmap of sterol proportions grouped according to B-ring saturation (cyclopropane ring (CPR), Δ0, Δ5, Δ7, Δ8, NA). All sterols which could not be assigned a name using reference materials were categorised as NA for B-ring saturation. In addition, ergosterol which is Δ5 and Δ7 was classed as NA. This heatmap shows a dominance of Δ5 sterols across the dataset and very low proportions of Δ0 sterols (stanols). Outer bars show the proportion and identity of the most dominant sterol in each species, showcasing the diversity of sterol production of pollen. All named sterols are grouped according to their position in the sterol synthesis pathway. *A)* Despite the diversity of the sterolome in many pollens, some species are strongly dominated by a single sterol. This can be seen in these Asparagaceae pollens which mostly contain schottenol. *B)* In some areas of the tree, sterols close together in the synthesis pathway are produced in high proportions by related species. In this case, β-sitosterol, avenasterol and isofucosterol are the dominant sterols in a range of Fabaceae, Salicaceae and Hypericaceae species. *C)* There are clear switches in the dominant sterol between some taxa; within the two Rosaceae subfamilies plotted, Amygdaloideae is dominated by 24-methylenecholesterol whilst Rosoideae is dominated by desmosterol. *D)* Asteraceae represented our most sampled family and showed an interesting sterol profile compared to other families. Species contained higher proportions of sterols with a cyclopropane ring or where B-ring saturation could not be determined (NA) than other families.

Typically, the pollen sterol profile was dominated by 1-3 major sterols (Supplementary - Figure 2). β-sitosterol and isofucosterol were the most common major sterol components (defined as ranking in top three by proportion of any given species), in over 100 taxonomic units (Supplementary - Table 4). Some plant species had a clear dominant sterol. Schottenol, 24-methylenecholesterol, desmosterol, cycloartenol and isofucosterol all occurred at >75% in at least one taxonomic unit (Supplementary - Table 1). In total, 29 species of the 261 which could be placed on our phylogeny had a sterol which accounted for ≥65% of their total. In 11 out of 29 species this sterol was 24-methylenecholesterol.

**Figure 2.**
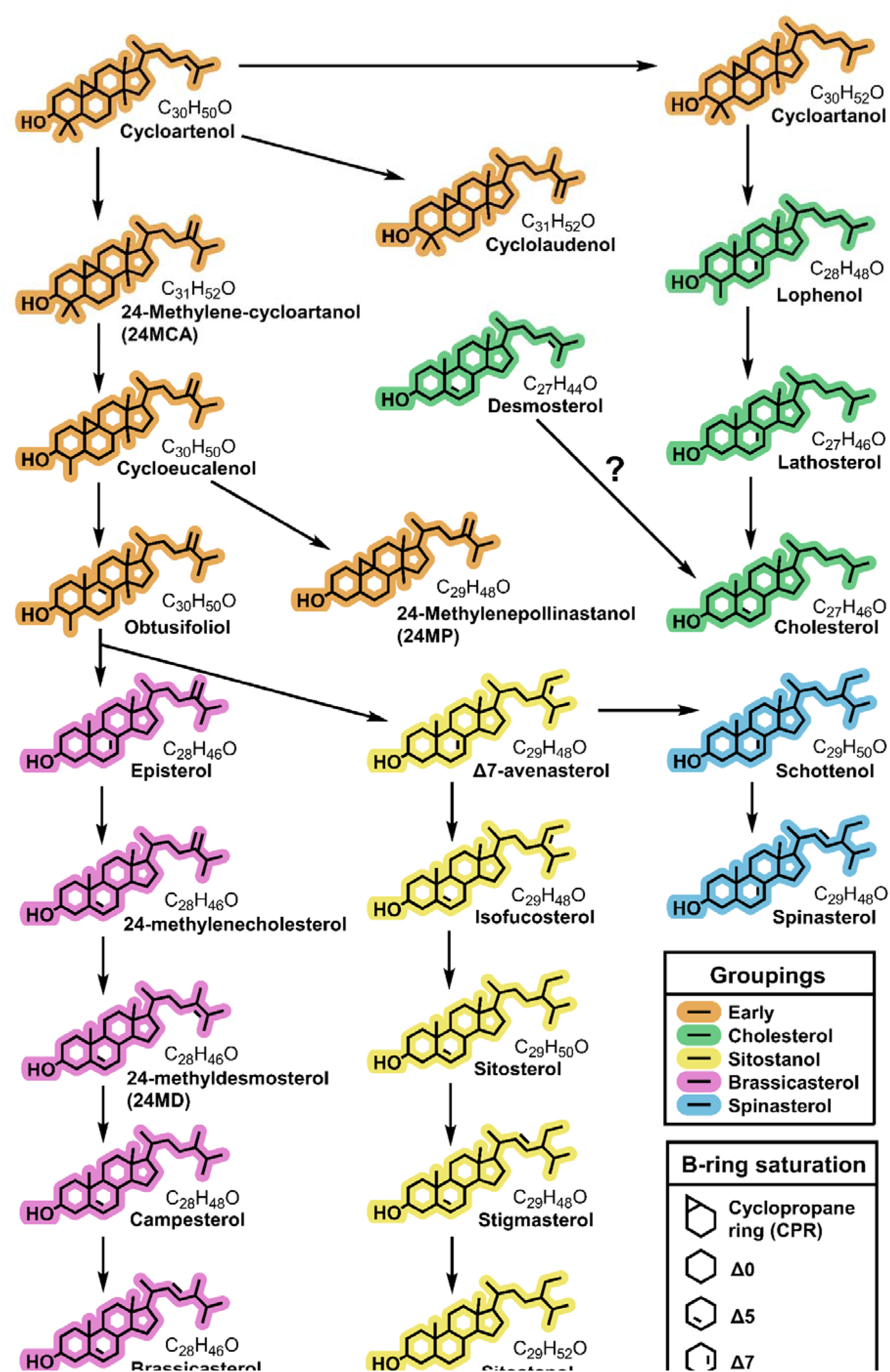
Sterol synthesis pathway in plants collated from (Sucrow and Radüchel, 1971; Schaller, 2003; Morikawa et al., 2006; Desmond and Gribaldo, 2009; Villette et al., 2015; Sonawane et al., 2016). All sterols which could be named in this study using reference materials are shown, excluding ergosterol which is not synthesised by plants. Sterols which could only be identified to level two annotation (carbon count and double bond saturation) are not shown. Sterols are coloured by section of the synthesis pathway to highlight structural similarities such as carbon count and emphasise branching points in the synthesis pathway. Many of the sterols abundant in pollen, such as 24-methylenecholesterol, isofucosterol and β-sitosterol, are not the end products of these synthesis pathways

Of the 69 sterols which occurred at >1% in pollen, 31 showed a significant phylogenetic signal for both metrics tested (K and λ), along with total sterol content (mg/kg) (Supplementary - Table 5). Furthermore, when sterols were grouped according to number of carbons (CC: 27-31) and B-ring double bond substitutions (DB: CPR, 0, 5, 7, 8, NA (*level 2 annotated sterol structures and ergosterol (*Δ*5&7*))), most groups showed significant phylogenetic signal for both metrics (K and λ) (27, 28, 29, 30 (CC) and CPR, 0, 5, 7, NA (DB)). Twenty-six additional sterols showed significant phylogenetic signal for a single metric (all Pagel’s λ except salisterol). The 31 carbon group and Δ8 group were also significant for Pagel’s λ.

Figure 1 presents the phylogenetic tree covering 261 of the 295 taxa analysed representing 54 families (a labelled species tree is provided in Supplementary - Figure 3). The dominant sterol produced in pollen was plotted across taxa. In this figure, for example, the four Asparagaceae species were dominated by a single sterol, schottenol, at proportions exceeding 70% (Fig 1A). Many species across Fabaceae, Salicaceae and Hypericaceae produced high levels of one of either isofucosterol, β-sitosterol or avenasterol (Figure 1B). These sterols arise from the same biosynthetic pathway and therefore might be expected to co-occur in related species (Figure 2). By contrast, in the closely related subfamilies of the Rosaceae, each exhibited a different sterol profile. The subfamily Rosoideae, was dominated by desmosterol whereas its sister subfamily, the Amygdaloideae, was dominated by 24-methylenecholesterol (Figure 1C). The sterolome for the Asteraceae was different from all other families in the tree. This family exhibited a consistently low proportion of Δ5 sterols, and instead produced pollen with high proportions of obtusifoliol or cycloeucalenol; sterols occurring early in the biosynthetic pathway (Figure 1D).

**Figure 3.**
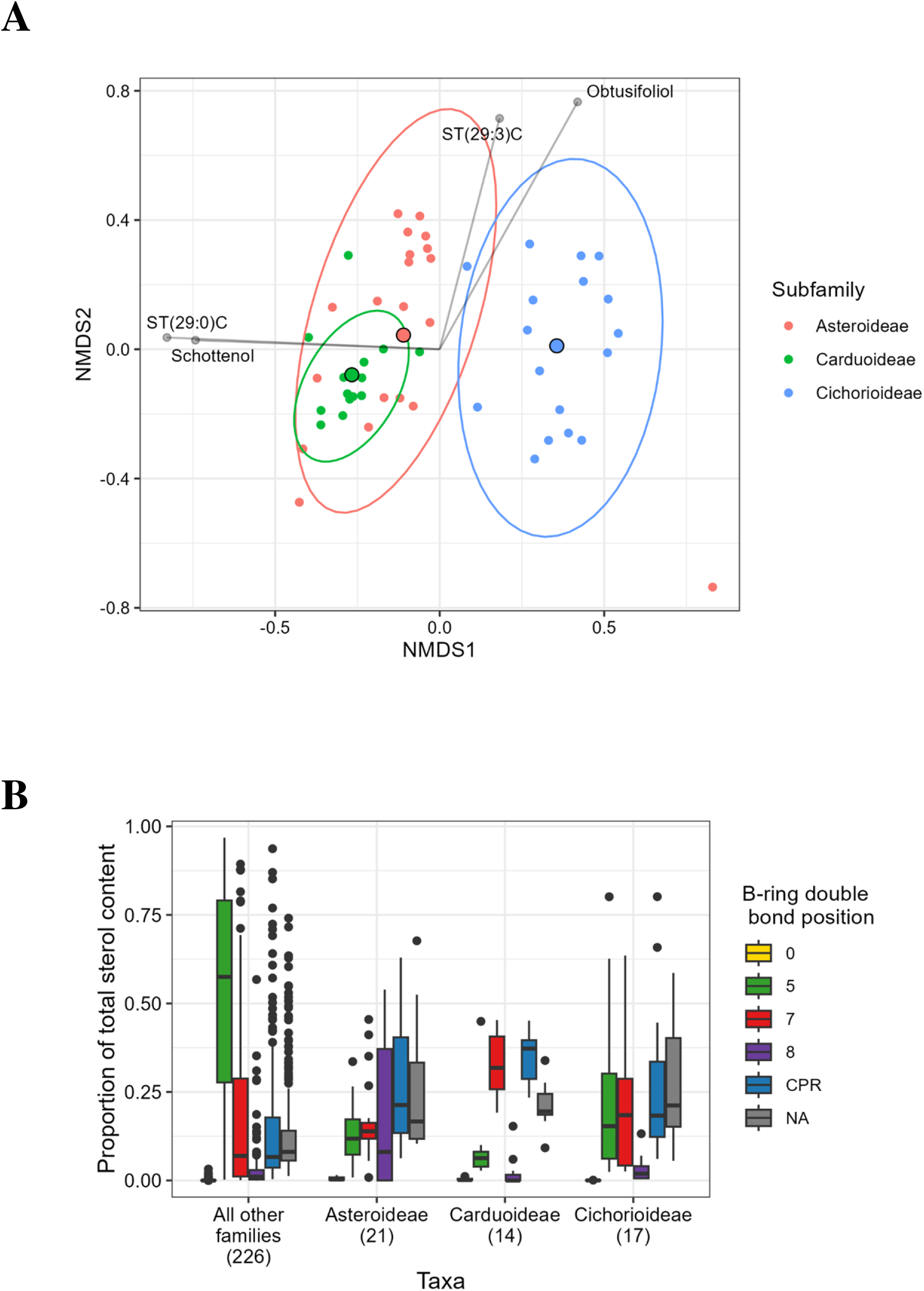
***A)*** NMDS biplot showing differences in sterol profile between Asteraceae subfamilies Asteroideae, Carduoideae and Cichorioideae. All pairwise differences between groups were significant (PERMANOVA, p<0.005) and each subfamily was associated with at least six level two annotated sterols which distinguished them from other Asteraceae species (ISA, Supplementary – Table 6). Plot stress value was 0.166 and groups showed significantly different variances with Carduoideae species showing the smallest variation in sterolome. Asteroideae and Carduoideae species also showed the greatest overlap in sterolome. ***B)*** Comparison of sterol composition between Asteraceae subfamilies (Asteroideae (21 spp.), Carduoideae (14 spp.) and Cichorioideae (17 spp.)) and all other species (226 spp.) when sterols are grouped by B-ring saturation (cyclopropane ring (CPR), Δ0, Δ5, Δ7, Δ8, NA, as shown in Figure 2). On average, non-Asteraceae species were dominated by Δ5 sterols. In contrast, Asteraceae flowers had higher levels of CPR, Δ7 and Δ8 sterols. Within Asteraceae, Carduoideae had a dominance of Δ7 sterols and Asteroideae showed a higher proportion of Δ8 sterols. Plot shows first and third quartile, median and 1.5 x inter quartile range.

### Phylogeny predicts profile: conservation of sterol biosynthesis at subfamily, genus, and species levels

There was a strong influence of phylogenetic group on the pollen sterolome, even at subfamily level. The sterol profiles of all three Asteraceae subfamilies differed significantly (PERMANOVA: F_(2,49)_ =10.234, *p*=0.001, R^2^=0.295, all pairwise comparisons *p*=0.001, comparison of variance: F_(2,49)_ =8.003, *p*<0.001) as shown in Figure 3A (NMDS plot stress value: 0.166). Our analysis indicated that each subfamily was associated with *at least* six level 2 annotated sterols. For example, the Carduoidae had 15 of these putatively identified sterols that distinguished it from the Asteroideae and the Cichorioideae (Supplementary - Table 6) indicating there is still much we do not know about sterol production in plants and especially in the Asteraceae.

Overall and when compared to other families, Asteraceae pollen had a highly distinctive sterolome (Figure 3B). There were also significant differences between Asteraceae subfamilies and other pollen in all sterol double bond classes (CPR: H(3)=49.6, *p*<0.001, Δ0: H(3)=78.6, *p<*0.001, Δ5: H(3)=55.8, *p<*0.001, Δ7: H(3)=17.0, *p<*0.001, Δ8: H(3)=14.2, *p<*0.005, ΔNA: H(3)=41.6, *p<*0.001). All Asteraceae subfamilies had significantly higher proportions of sterols with a B-ring cyclopropane substitution (CPR), undetermined B-ring structures (ΔNA – Level 2 annotated sterols), and lower proportions of Δ5 sterols than all other families (*post-hoc* Dunn tests: CPR: *p*<0.005, Δ0: *p<*0.001, Δ5: *p<*0.005, ΔNA: *p<*0.001). Furthermore, Carduoideae had significantly higher Δ7 sterols than all non-Asteraceae families (*p<*0.001) and significantly lower Δ8 than all other groups (other families: *p<*0.010, Asteroideae: *p<*0.005, Cichorioideae: *p<*0.005).

Comparing between genera of the same family we found significant differences in sterolome, demonstrating that species within a genus often display similar sterolomes (Figure 4A). In Asteraceae, *Centaurea* and *Cirsium* both belong to the subfamily Carduoideae and show more similar sterolomes whilst *Crepis* belongs to the Cichorioideae. However, genera were still significantly different from each other (PERMANOVA: Genus: F_2,20_=26.087, *p*=0.001, Species: F_9,20_ = 2.311, *p*<0.010. Variance: Genus: F_2,29_=0.810, *p* >0.100, Species: F_11,20_=1.543, *p*>0.100). All pairwise differences between genera were significantly different (all *p*=0.001) with genus and species identity together explaining a high proportion of the variance in the dataset (R^2^: genus = 0.561, species= 0.224). All genera were associated with at least five level two annotated sterols, affirming findings above about the undetermined sterol composition of Asteraceae flowers (Supplementary - Table 7). This pattern was repeated in the three Fabaceae genera *Trifolium*, *Ulex* and *Vicia* (PERMANOVA: Genus: F_2,22_=28.756, *p*=0.001, Species: F_8,22_ = 2.567, *p*=0.001. Variance: Genus: F_2,30_=1.212, *p*>0.100, Species: F_10,22_=2.423, *p*<0.05, Pairwise differences: all *p*<0.005, R^2^: Genus = 0.575, Species = 0.205). The strongest associations between genera and sterols were *Trifolium* with schottenol and *Ulex* with sitostanol. In the Rosaceae, strong differences were recorded between genera, even within the same subfamily, however, species-level differences within genera were not statistically significant, despite *Potentilla* and *Rubus* both belonging to the same subfamily Rosoideae (PERMANOVA: Genus: F_2,17_=16.695, *p*=0.001, R^2^ = 0.632, Species: F_6,17_ = 0.414, p>0.500, R^2^ = 0.047. Variance: Genus: F_2,23_=1.183, *p*>0.100, Species: F_8,17_=0.417, *p>*0.500). Pairwise comparisons: *Prunus*: *Potentilla* (*p*<0.005), *Prunus*: *Rubus* (*p*<0.005), *Potentilla*: *Rubus* (*p*<0.050). All three genera were associated with a high number of level two annotated sterols but the strongest association was *Rubus* with desmosterol.

**Figure 4.**
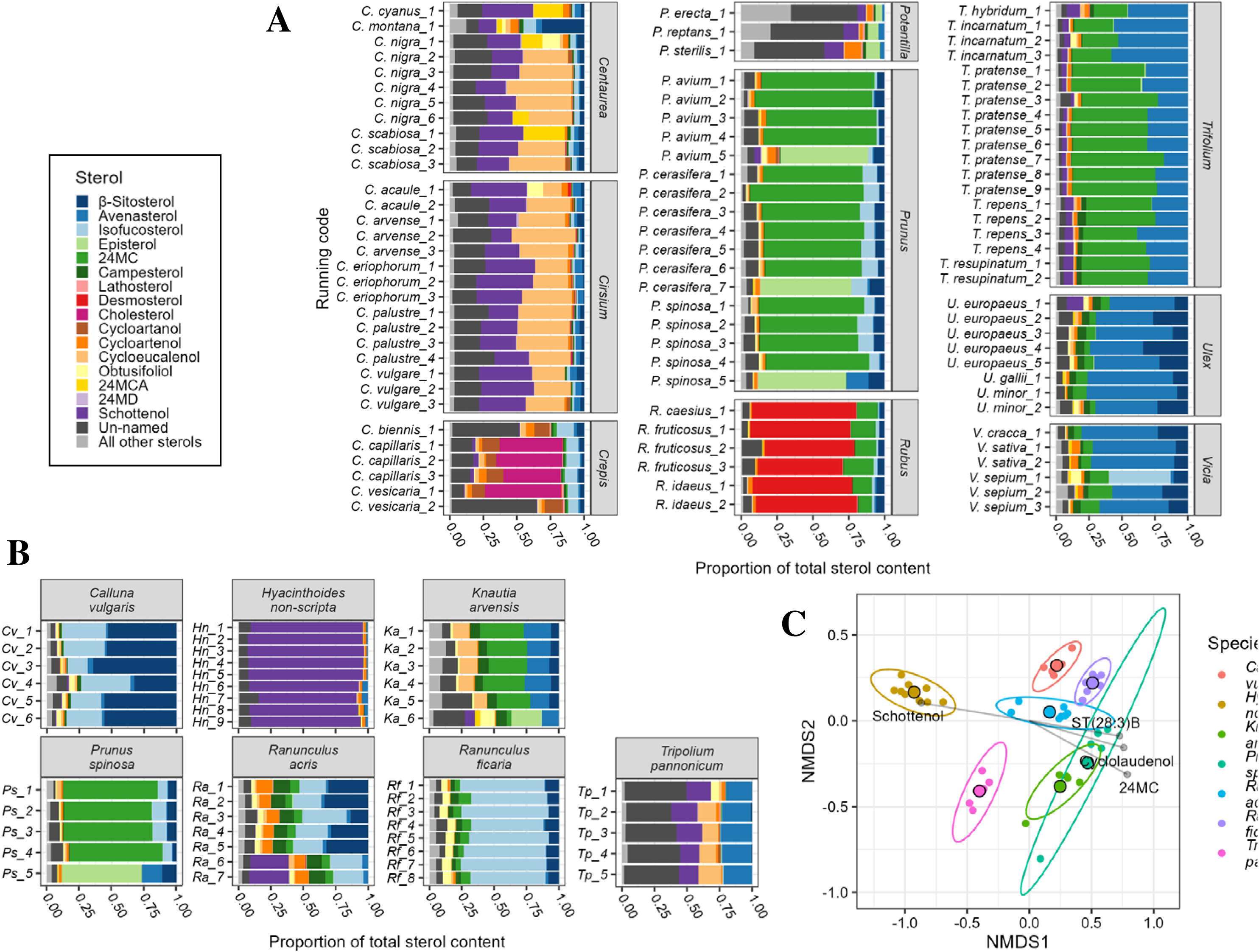
***A)*** Sterol composition of species belonging to repeatedly sampled genera in the most sampled families; Asteraceae, Fabaceae and Rosaceae. The magnitude of inter-genus differences appears to vary widely; some confamilial genera share similar sterolomes (Vicia and Ulex, Centaurea and Cirsium) whilst others are dominated by different sterols (Rubus and Prunus, Crepis and Cirsium). Within genera, sterolomes appeared largely consistent between species. All non-major sterols have been collapsed into ‘All other sterols’ and all sterols annotated to level two (carbon count and double bond saturation) are included as ‘Un-named’. ***B)*** Sterol composition of species collected from at least four counties and with at least five samples total. Tripolium pannonicum shows the highest proportion of sterols which could only be annotated to level two (carbon count and double bond saturation) suggesting there is still much we do not know about sterol production in this species. Species generally show a strong consistency in sterolome, this is reflected in tight group distributions when plotted by NMDS (Figure 4C). All non-major sterols have been collapsed into ‘All other sterols’ and all sterols annotated to level two are included as ‘Un-named’ ***C)*** NMDS plot of proportion data from samples shown in Figure 4B arcsine transformed. Difference among species groups were significant (p=0.001) and variance was not significantly different (p= 0.729). This suggests there is consistency in sterol profile between variable abiotic conditions which aligns with a strong genetic control of sterol production, as indicated by the phylogenetic signal across the data. Lines show associations with different sterols, only sterols with significant (p<0.0100 and strong >0.7) associations were used. Stress value of NMDS was 0.096.

Furthermore, we examined variation in the sterol profile of individual species collected in different sites across the UK. Intraspecies variation between sites was not greater than the interspecies variation in the dataset. Species displayed significantly different sterol profiles (PERMANOVA, F_6,39_ = 33.757, *p=*0.001, all pairwise comparisons between species *p<*0.050), with species identity explaining a large proportion of the variance in the data (R^2^=0.839). In contrast, the sterol profile of pollen collected from the same species in different locations across the UK exhibited relatively little variation (Figure 4B, C (stress value = 0.096)). Interestingly, the amount of variation in the sterol profile within species was not significantly different (F_6,39_ = 0.600, *p>*0.500). The sea aster, *Tripolium pannonicum,* was significantly associated with the most individual sterols (14), including a very strong association with ST(27:1)B. The meadow buttercup, *Ranunculus acris,* had a strong association with ergosterol. Since this sterol is produced exclusively by fungi it suggests buttercups are more likely to be contaminated with fungi. Species that collect pollen from this species are therefore likely to be provisioning significant quantities of fungi to their brood. The common bluebell, *Hyacinthoides non-scripta,* was strongly associated with schottenol, the lesser celandine, *Ranunculus ficaria,* with isofucosterol, blackthorn (*Prunus spinosa)* with 24-methylenecholesterol and heather (*Calluna vulgaris)* with sitostanol, β-sitosterol and stigmasterol. A full list of all 45 associations found can be seen in Supplementary Table 8.

### Comparison of sterols found in honeybees and pollen

The honeybee sterol profile from Svoboda *et al*., (1980) was compared to the sterol profile of all hand collected pollen species in this dataset (Figure 5). Bee sterols differed significantly from pollen in sterol profile (PERMANOVA: F_(1,294)_ = 10.567, *p*=0.001, R^2^=0.035) and in variance in sterol profile (F_(1,294)_ = 98.678, *p*<0.001). The pollen sterolomes that were most like honeybees were replotted from the full NMDS (stress=0.101) in Figure 5A and their sterol profiles shown in Figure 5B.

**Figure 5.**
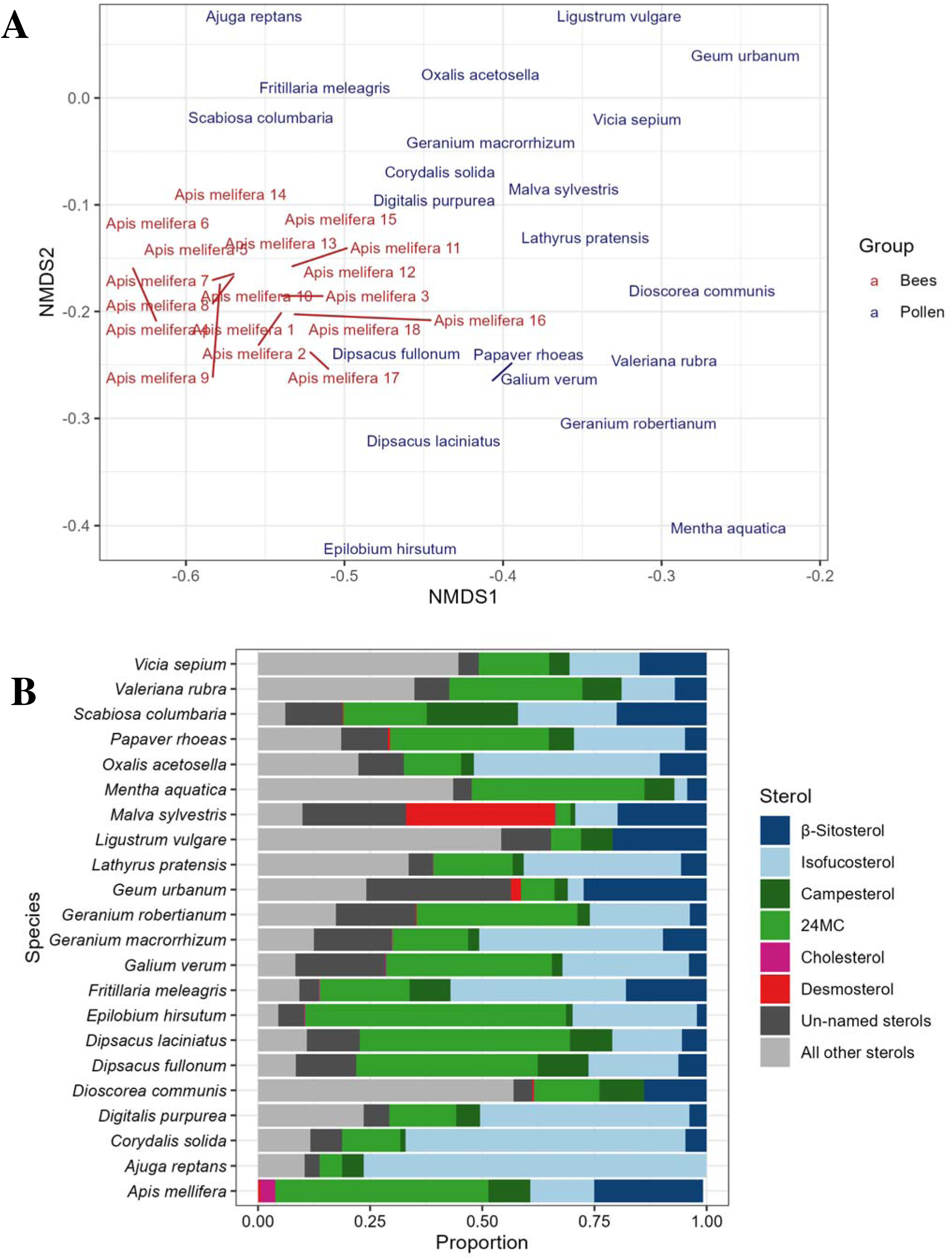
*A)* Subsection of an NMDS comparing 18 honeybee prepupae sterol samples from Svoboda, et al., (1980) with all hand collected pollen units reported in this paper (278). Bees and pollen showed significantly different sterol profiles and the stress value of the 3D NMDS was 0.101. Only those pollens closest to the bees in NMDS space are plotted here in 2D for clarity. Many species that are popular with honeybees, for instance Rubus fruticosus, did not cluster closely with honeybees suggesting that similarity in sterol profile alone does not motivate foraging by honeybees. Pollens showed a much wider variation in sterolome than the bees though this is to be expected as they cover a much wider ecological and taxonomic range. Pollen also contains a much wider array of sterols compared to honeybees and so no pollen is likely to directly resemble honeybee tissues. ***B)*** Proportions of honeybee-relevant sterols for all plant species shown in Figure 5A plus a mean of the 18 honeybee prepupae sterol samples from (Svoboda et al., 1980). All pollens contained high proportions of 24-methylenecholsterol, β-sitosterol and isofucosterol. However, many pollens contained more isofucosterol and less 24-methylenecholesterol and β-sitosterol than bees. In addition, honeybee prepupae contained higher proportions of cholesterol than any of the similar pollen. It is again clear that no individual pollen will provide all the sterols required by honeybees in their requisite proportions.

We used data from Svoboda *et al*., (1980)’s analysis of prepupal honeybees to estimate the sterols required by bee species although the sterol requirements of other bee species may differ from this. From these data, we estimated that bees required >30% of 24-methylenecholesterol, >19% of β-sitosterol, >10% of isofucosterol, >5% of campesterol, >0.5% of cholesterol and > 0.5% of desmosterol. Using these thresholds, we found that 88 species did not meet any of the sterol requirements of honeybees (Figure 6A). Strikingly, no species met all the required proportions of all six sterols, even though 188 species contained all six sterols. Of these sterols, 24-methylenecholesterol has been argued to be the most important (Svoboda et al. 1980, others); only 35 out of 278 taxonomic units contained sufficient 24-methylenecholesterol to meet this threshold (Figure 6A). Out of these 35, only 12 also contained enough isofucosterol; none of these 35 contained enough β-sitosterol. Other species were sufficient in individual bee-relevant sterols. For example, 82 species contained enough β-sitosterol and 90 contained enough isofucosterol: 48 contained enough of both compounds. Likewise, 62 contained enough campesterol. The fewest species contained sufficient cholesterol (21) or desmosterol (28).

**Figure 6.**
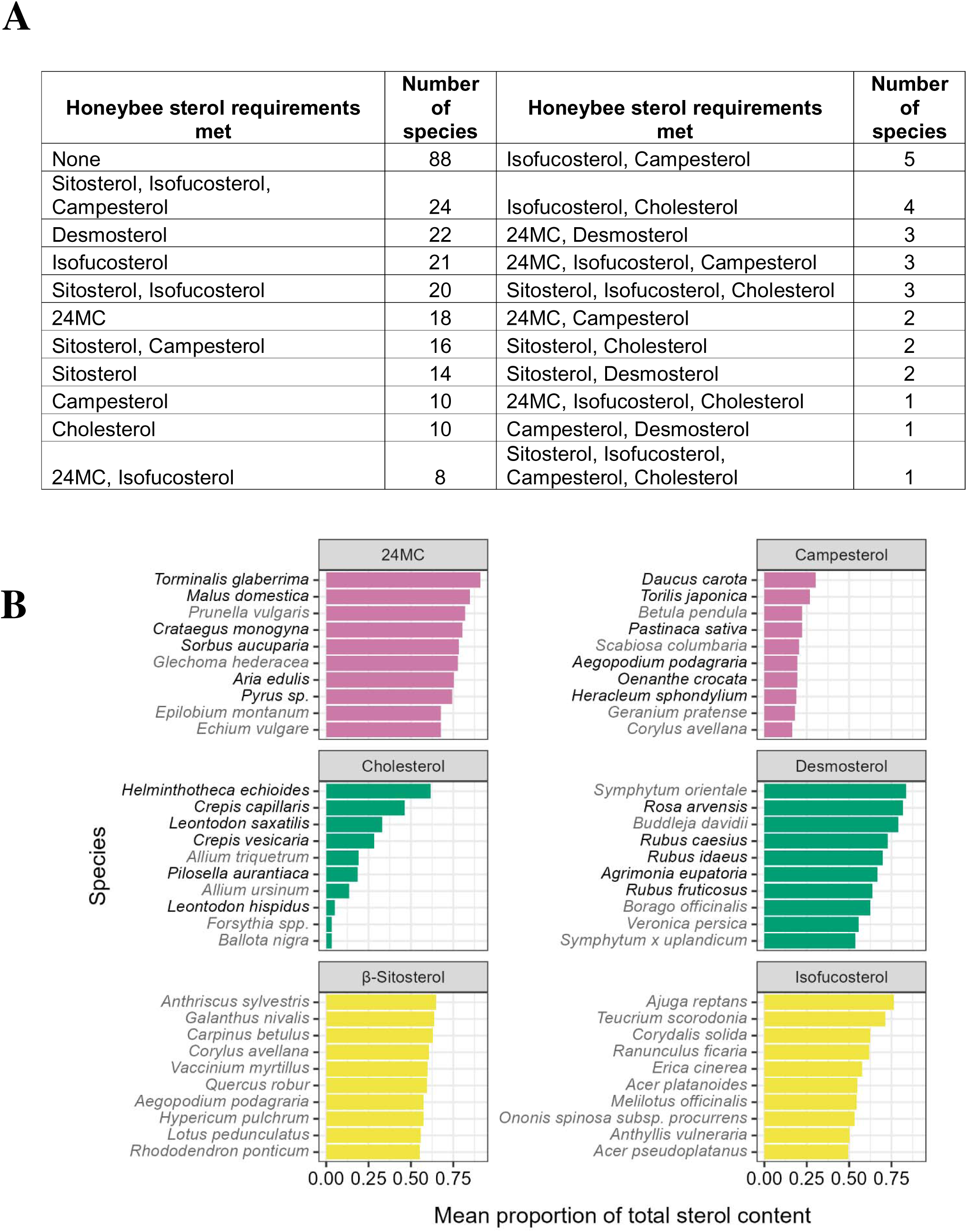
***A)*** Summary of how many pollen species met the sterol requirements identified from analysis of prepupal honeybee tissues (24-methylenecholesterol >30%, β-sitosterol >19%, isofucosterol >10%, campesterol >5%, cholesterol >0.5%, desmosterol >0.5% - Svoboda et al., (1980)). Out of 278 species, 88 met none of these requirements, only 32 contained three or more at the required levels. A single species, Ballota nigra, fulfilled four of the six required (β-sitosterol, isofucosterol, campesterol and cholesterol). ***B)*** The ten highest producing species for the six sterols detected in honey bee tissues (Svoboda et al., 1980). Campesterol and cholesterol are only ever present in lower proportions and are never dominant in a pollen, unlike 24-methyelenecholesterol, desmosterol and isofucosterol which are over 75% of total sterols in the highest producing species. Similarities in sterol production between related species mean many of the top species for a sterol belong to a single family. Dominant families are highlighted by species names in black. 24-methylenecholesterol: Rosaceae, Campesterol: Apiaceae, Cholesterol: Asteraceae, Desmosterol: Rosaceae. Sterols are coloured according to their positioning in the plant sterol synthesis pathway in Figure 2.

The highest proportions of the honeybee relevant sterols were often found in specific plant families as shown in Figure 6B (24-methylenecholesterol = Rosaceae, campesterol = Apiaceae, cholesterol = Asteraceae subfamily Cichorioideae). Interestingly, the highest content of desmosterol was from diverse flowers popular with honeybees and bumblebees such as *Rubus* spp., *Rosa* spp., *Borago officinalis* and *Symphytum* sp. β-sitosterol was not recorded more frequently by any family and three of the species in which this sterol was most abundant are wind pollinated trees (*Carpinus betulus*, *Corylus avellana*, *Quercus robur*).

## Discussion

This dataset provides a major contribution to our knowledge of pollinator nutrition at a scale never previously undertaken and informs our understanding of the evolution of pollen sterol pathways across angiosperms. We also show that abiotic factors do not significantly influence sterol profiles and that instead they are under evolutionary constraints.

Sterols are needed for pollen tube elongation. However, these sterols can be synthesised *de novo* from squalene during pollen tube growth (Villette *et al*., 2015). It is possible that storage of sterols in pollen grains would make them a target for insect herbivores, which could have a dramatic negative effect on plant fitness. The high diversity of uncommon sterols in pollen could therefore be a mechanism for the plant to escape pollinivory, as a diverse sterolome could frustrate herbivore adaptation to targeted efficient sterol conversion pathways. It is possible that such sterols could either be unusable or even toxic to pollinivores. During their evolution, pollinivores including bees may have adapted to be able to use the diverse range of pollen sterols to overcome this defence. Previous studies have hypothesized that bees had lost the genes for the dealkylation of longer chain sterols to cholesterol and instead adapted a specific subset of pollen sterols for their use (Svoboda, Herbert and Thompson, 1983). This would also allow bees to use several different sterols flexibly rather than relying on obtaining a single sterol abundant enough for them to convert efficiently. Overcoming such a defence would also enable bees to use a valuable nutritional resource unavailable to other pollinivores.

The sterols most abundant in honeybees (24-methylenecholesterol, β-sitosterol and isofucosterol) are also commonly abundant in pollen sterolomes. The structural properties of these sterols (C28:Δ5, C29:Δ5 and C29:Δ5 respectively) may be useful in bee cell membranes, influencing properties such as fluidity (Furse, Koch, *et al*., 2023). The dominance of these nutrients in honeybees may represent an adaptation of bees to use the sterols available in pollens. It is interesting that only 29 out of 261 species in the phylogeny had a sterol which accounted for ≥65% of total sterols. Ten out of 29 of these species were Rosaceae with the dominant sterol being 24-methylenecholesterol or desmosterol. In fact, 24-methylenecholesterol was the most common sterol that accounted for ≥65% of total sterols (11 species).

It is conceivable that co-evolution of plants and bees has been driven in part by the specific sterols in pollen to encourage bee visitation and support bee populations although this is more likely the case in specialist bees (Vanderplanck et al., 2017). However, the six sterols required by honey bees (24-methylenecholesterol, β-sitosterol, isofucosterol, campesterol, cholesterol, desmosterol) were not recorded in the pollen of any one single plant species in exactly the same proportions recorded in bees. Notwithstanding that floral constancy presents benefits to plants of pollen being delivered to conspecifics, it is also conceivable that by producing sterols in quantities that are less than ideal for a pollen feeder that this could potentially prevent overharvesting of pollen from one source and instead encourage a generalist foraging strategy. The miss-match between bee nutritional needs and pollen chemistry (especially among crops such as *Malus* spp.) may help explain the effectiveness of agri-environmental floral plantings in improving crop pollination. It is possible that where sterol requirements are known that such plantings might be specifically selected for complementary nutritional properties.

It is plausible that sterol diversity in honeybees is a result of their generalised diet and their nutritional flexibility. However, the pollen sterolome also contains a range of sterols that are absent from the tissues of honeybees such as obtusifoliol, schottenol and cycloeucalenol. While these sterols may serve key physiological functions for plants unlike Δ5 sterols they may be unsuitable as replacements for cholesterol in insect tissues (e.g., sterols with B-ring cyclopropane substitutions or Δ8 double bond).

The ‘Asteraceae paradox’ refers to the frequent use of Asteraceae pollen by specialists and its infrequent collection by generalists (Müller and Kuhlmann, 2008). It is thought to be the result of chemical defences and nutritional deficiencies (Praz, Müller and Dorn, 2008; Vanderplanck, Gilles, *et al*., 2020). Our results showed that Asteraceae pollen contained lower proportions of Δ5 sterols such as 24-methylenecholesterol, β-sitosterol, isofucosterol desmosterol and campesterol – typically favoured by honeybees, instead producing higher quantities of Δ7, Δ8, un-named (L2) or those sterols with B-ring cyclopropane substitutions. This supports the hypothesis that Asteraceae pollen is nutritionally less suitable for species of bees that typically forage for pollen which contains high levels of Δ5 sterols such as honey bees. Furthermore, sterols with B-ring cyclopropane substitutions occur early in the sterol synthesis pathway and are therefore energetically less costly to produce than sterols which require additional metabolic steps. The Asteraceae sterolome may therefore represent a strategy to escape pollinivory by producing un-useable sterols which further benefit the flower by acting as precursors for the synthesis of downstream sterols during pollen tube growth. Additionally, the low proportions of Δ5 sterols in Asteraceae flowers suggests the sterol requirements of Asteraceae-specialist solitary bees may have evolved different phytosterol use to that of common eusocial species (*Apis* spp. and *Bombus* spp.) and provides more support to the nutritional deficit argument of the Asteraceae paradox.

The variation in both sterol profile and total sterol content between species leads to a stark difference in sterol availability between flowering plants. When considering the availability of pollen nutrients in floral seed mixes, it is important to consider their accessibility to different pollinators through floral morphology and phenology. The strong phylogenetic signal and within-genus consistency in sterol profile presented here invites further investigation into the mechanisms regulating sterol production in pollen from an evolutionary perspective. An assessment of stigma-ovule distance in the study species may prove enlightening for the trends in total sterol production.

## Acknowledgements

We would like to acknowledge the organisations who granted access for sample collection: Christchurch College gardens, Wytham Woods, Oxford City Council, Natural England, Royal Botanic Gardens, Kew, Warwick District Council, The University of Oxford Botanic Garden & Arboretum, The University of Oxford University Parks, FAI farms, Friends of Aston’s Eyot, The Wildlife Trusts, Hogacre Common Eco Park, St Johns College, Salisbury Plain Training Area (SPTA) Conservation Group and the Ministry of Defence, The Butterfly Conservation Trust, The National Trust and Shotover Preservation Society. We also acknowledge PlantLife for their support of the project. We thank Samuel Furse and Geoff Kite in the biochemistry laboratory at Kew for support providing the sterol data and Hauke Koch for his early contributions to the study. We also thank Judy Webb for her help with plant identification and to Daniel Stabler and Jennifer Chennells for their help in method development. Sample collection was assisted by Flynn Bizzell, Harry Savage, Nell Miles, Poppy Emson and Léa Beaupère. This work was supported by a NERC-UKRI grant NE/V012282/1 to PCS and GAW, a BBSRC grant BB/T014210/1 to PCS, a BBSRC grant BB/T015292/1 to GAW and NERC-UKRI PhD funding (NE/S007474/1) through the Oxford DTP in Environmental Research to ECB.

