## SUPPLEMENTARY FILES for "Pollen sterols are highly diverse but phylogenetically conserved"

***Table 1.*** *Maximum, minimum, median and Inter Quartile Range (IQR) percentage of sterols which occurred at >40% in at least one species. Statistics were calculated from species/subspecies means, (genus where species was not available). Sterols which have a median of >2.5%, isofucosterol, avenasterol and β-sitosterol, are highlighted in grey. These sterols are positioned close together in the sterol biosynthesis pathway (Figure 2). For un-named sterols, a level two annotation (carbon count and double bond saturation) is shown in brackets.*

| **Sterol** | **Max** | **Min** | **Median** | **IQR** | **Percentage of species detected in** |
| --- | --- | --- | --- | --- | --- |
| 24MC | 91.03 | 0.00 | 1.51 | 11.47 | 99.3 |
| Schottenol | 88.24 | 0.00 | 0.25 | 5.04 | 99.3 |
| Desmosterol | 83.50 | 0.00 | 0.02 | 0.05 | 91.5 |
| Cycloartenol | 77.92 | 0.02 | 2.32 | 4.25 | 100.0 |
| Isofucosterol | 76.41 | 0.00 | 3.47 | 14.64 | 98.0 |
| ST(29:1)C | 70.94 | 0.00 | 0.13 | 0.33 | 98.0 |
| Avenasterol | 69.79 | 0.00 | 2.58 | 12.95 | 99.3 |
| ST(30:2)B | 65.65 | 0.02 | 0.22 | 0.48 | 100.0 |
| β-Sitosterol | 64.90 | 0.00 | 4.68 | 20.24 | 96.3 |
| Cycloartanol | 62.49 | 0.00 | 0.03 | 0.10 | 91.9 |
| Episterol | 61.76 | 0.00 | 0.02 | 0.14 | 69.8 |
| Cholesterol | 61.67 | 0.00 | 0.01 | 0.11 | 73.2 |
| Cycloeucalenol | 60.85 | 0.00 | 1.67 | 4.77 | 93.6 |
| ST(27:2)D | 60.06 | 0.00 | 0.00 | 0.02 | 79.0 |
| ST(27:1)B | 59.69 | 0.00 | 0.00 | 0.00 | 50.2 |
| Obtusifoliol | 56.75 | 0.00 | 1.11 | 3.05 | 88.1 |
| Lathosterol | 55.95 | 0.00 | 0.06 | 0.29 | 98.0 |
| ST(30:1) | 49.62 | 0.00 | 0.02 | 0.15 | 87.5 |
| 24MD | 45.24 | 0.00 | 0.07 | 0.31 | 94.9 |
| ST(30:2)A | 44.07 | 0.00 | 0.19 | 0.24 | 99.7 |
| ST(30:2)E | 40.37 | 0.00 | 0.50 | 1.25 | 99.7 |

***Table 2.*** *Maximum, minimum, median and Inter Quartile Range (IQR) of sterols which were present in all taxonomic units analysed (minimum species value in dataset >0). Statistics were calculated from species means. For un-named sterols, carbon count and saturation are shown in brackets. Only four sterols out of 78 were detected in all taxonomic units. The two which could be named using reference materials do not share structural similarities and are many multiple metabolic steps apart in the sterol synthesis pathway.*

| **Sterol** | **Max (%)** | **Min (%)** | **Median (%)** | **IQR (%)** |
| --- | --- | --- | --- | --- |
| Cycloartenol | 77.92 | 0.02 | 2.32 | 4.25 |
| ST(30:2)B | 65.65 | 0.02 | 0.22 | 0.48 |
| ST(30:2)C | 36.51 | 0.04 | 0.74 | 1.15 |
| Campesterol | 30.23 | 0.01 | 1.78 | 3.90 |

***Table 3.*** *Highest and lowest Simpson’s diversity index values across all pollen species. Highest values are shown in the left of the table and lowest values are in the right. These values demonstrate that even though pollen contains a wide range of sterols, they do not generally produce them in equal proportions.*

| **Species** | **Simpson's diversity index** | **Species** | **Simpson's diversity index** |
| --- | --- | --- | --- |
| *Papaver cambricum* | 0.913 | *Torminalis glaberrima* | 0.169 |
| *Hypochaeris radicata* | 0.911 | *Malus sp.* | 0.204 |
| *Leontodon sp.* | 0.906 | *Muscari armeniacum* | 0.219 |
| *Campanula latifolia* | 0.904 | *Hyacinthoides non-scripta* | 0.262 |
| *Hedera helix* | 0.902 | *Hyacinthoides x massartiana* | 0.273 |
| *Anemonoides nemorosa* | 0.889 | *Malus domestica* | 0.273 |
| *Primula vulgaris* | 0.886 | *Symphytum orientale* | 0.295 |
| *Pulicaria dysenterica* | 0.886 | *Rosa arvensis* | 0.317 |
| *Geum urbanum* | 0.885 | *Prunella vulgaris* | 0.323 |
| *Phacelia tanacetifolia* | 0.884 | *Crataegus monogyna* | 0.347 |

***Figure 1.*** *Sterol compositions of the species with the highest (left) and lowest (right) Simpson’s diversity index as shown in Supplementary Table 3. Species with a low diversity index are generally dominated by a single sterol, compared to species with a higher diversity index which produce smaller proportions of a wider range of sterols. Species are arranged alphabetically.*

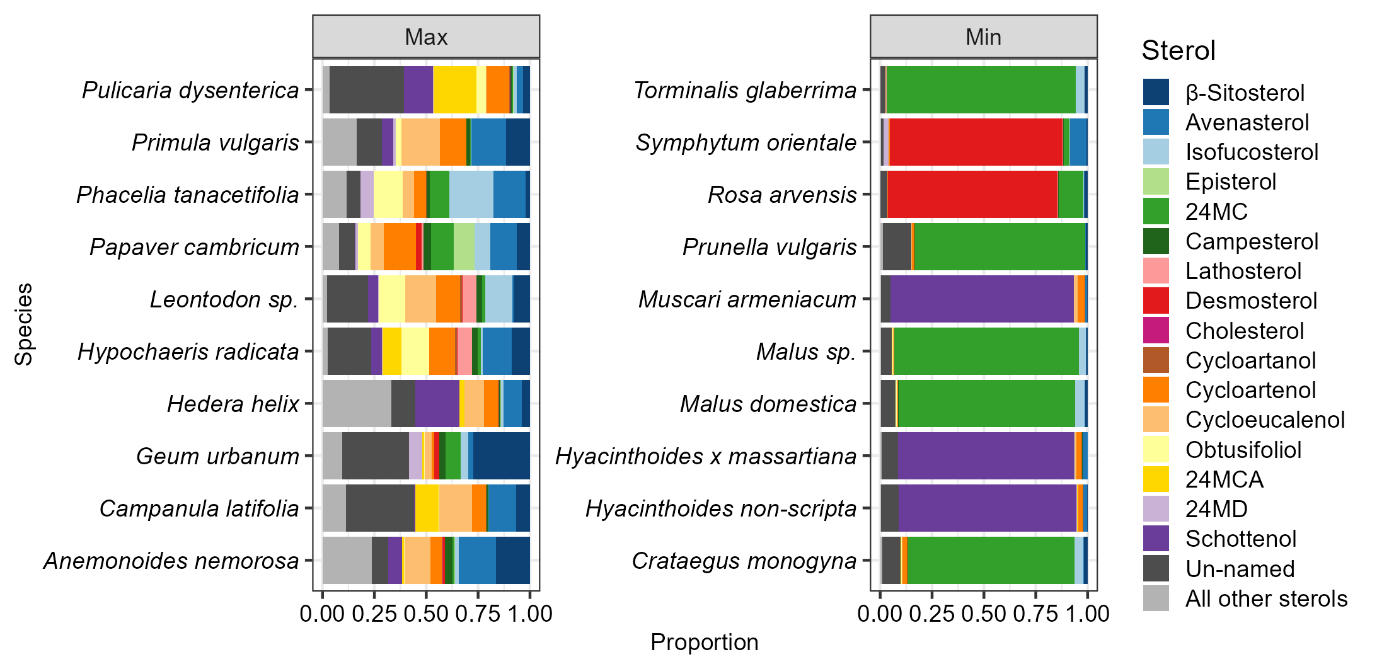

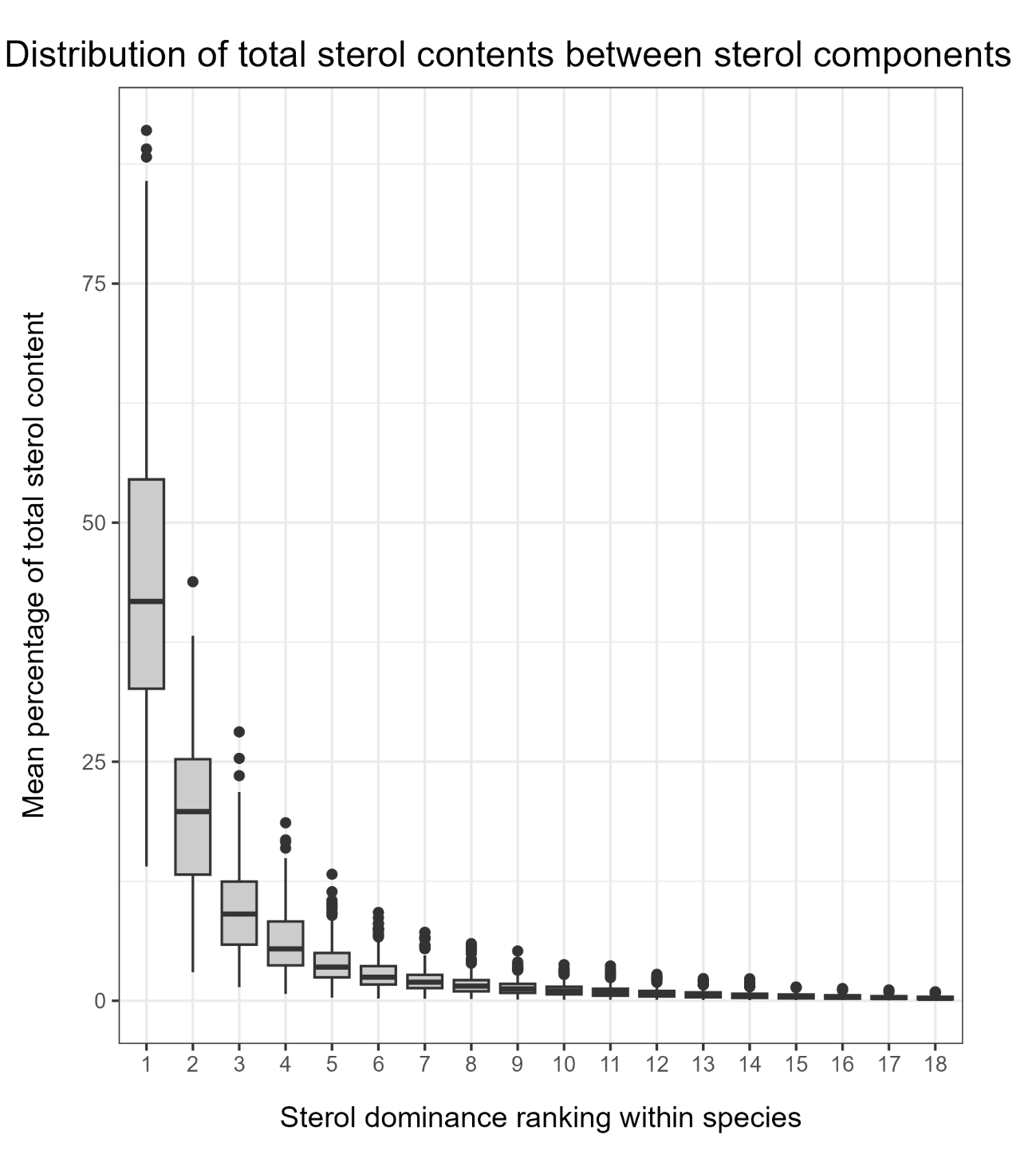

***Figure 2.*** *Median percentage and Inter Quartile Range (IQR) of the most dominant 18 sterol components within each taxonomic unit (295), ranked by proportion in the sample. For all taxonomic units, despite most containing >60 sterols, the most dominant 18 sterols accounted for at least 90% of total sterol content. The identity of these 18 sterols varied among taxonomic units.*

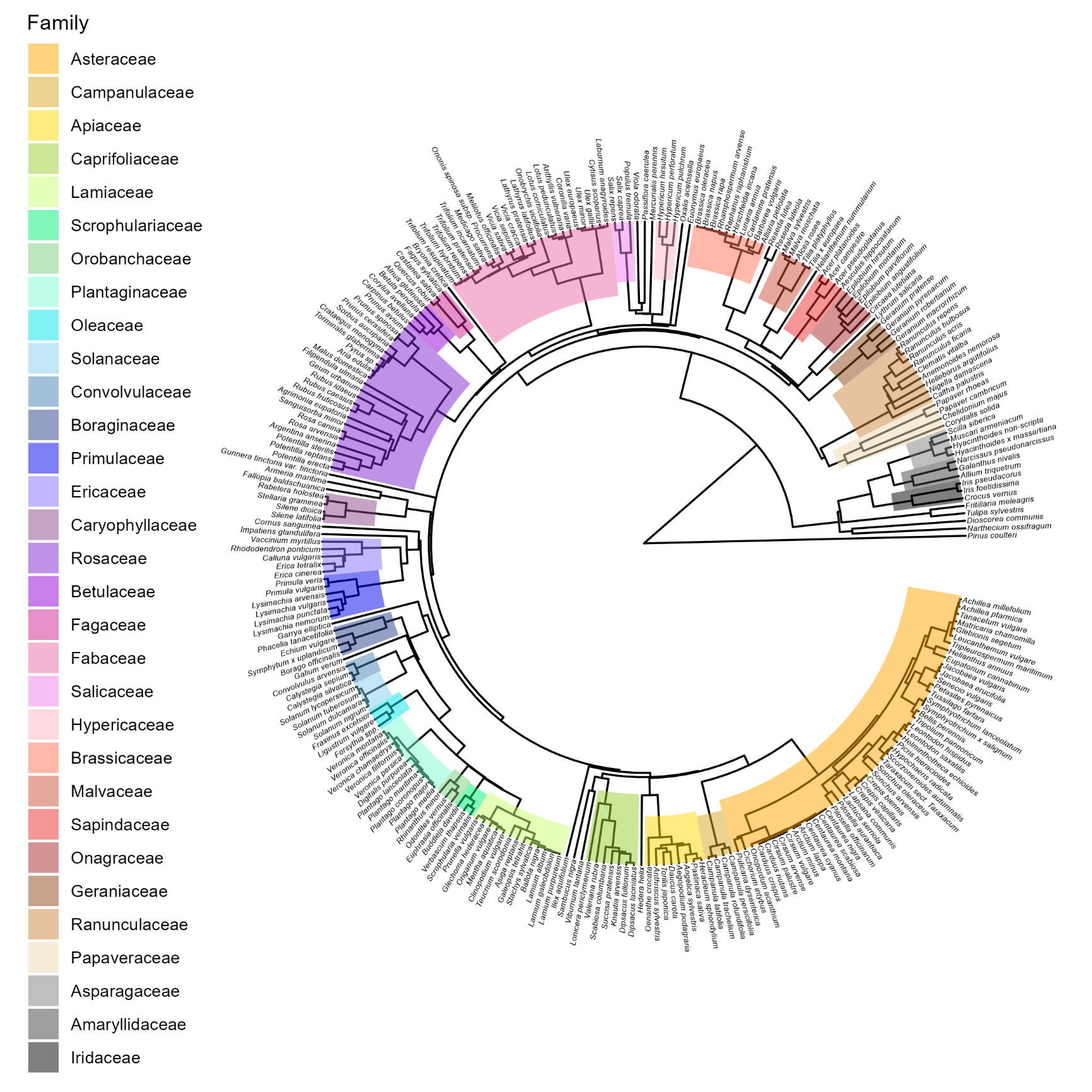

***Figure 3.*** *Labelled phylogeny shown in Figure 1 of main text, covering 261 species. Adapted from (Zanne et al., 2014). Families with three or more species are highlighted in a clockwise direction.*

***Table 4.*** *Sterols most commonly dominant in pollen, defined as occurring within the top three sterols by proportion. Sterols are ordered by number of taxonomic units for they were dominant in. Only two sterols were ranked in the top three for over 100 species; β-sitosterol and isofucosterol. Only 39/78 detected sterols ever occurred in the top three for any pollen species.*

| **Sterol** | **No. of species ranked in top three by proportion** | **Sterol** | **No. of species ranked in top three by proportion** |
| --- | --- | --- | --- |
| β-Sitosterol | 127 | ST(28:3)B | 9 |
| Isofucosterol | 120 | 24MCA | 8 |
| Avenasterol | 98 | ST(28:1)A | 8 |
| 24MC | 89 | ST(30:2)C | 8 |
| Schottenol | 61 | Cholesterol | 7 |
| Cycloeucalenol | 55 | ST(27:1)B | 3 |
| Campesterol | 46 | ST(27:2)E | 3 |
| Cycloartenol | 44 | ST(29:1)C | 3 |
| Obtusifoliol | 38 | Brassicasterol | 2 |
| ST(30:2)E | 23 | ST(27:1)D | 2 |
| Desmosterol | 20 | ST(30:1) | 2 |
| Cyclolaudenol | 13 | ST(31:2)A | 2 |
| ST(30:2)B | 13 | ST(31:2)B | 2 |
| Episterol | 12 | ST(28:3)A | 1 |
| 24MD | 11 | ST(29:0)C | 1 |
| ST(30:2)F | 11 | ST(30:2)A | 1 |
| 24MP | 10 | ST(30:2)D | 1 |
| Cycloartanol | 10 | Salisterol | 1 |
| Lathosterol | 10 | Stigmasterol | 1 |
| ST(27:2)D | 9 |  |  |

***Table 5.*** *Results of phylogenetic signal analysis on the proportions of all detected sterols with a maximum proportion >1% (69), total sterols (mg/kg), proportion of different carbon length sterols (light grey: 27, 28, 29, 30, 31) and proportion of different B-ring saturated sterols (dark grey: Cyclopropane ring (CPR), Δ0, Δ5, Δ7, Δ8, ΔNA). Only variables which were significant for both Bloomberg’s K and Paegel’s λ are shown. This includes 31/69 tested sterols and most structural groups, demonstrating the strong phylogenetic signal in the data. Ergosterol was omitted from this summary despite its significant phylogenetic signal as it is not part of the sterol synthesis pathway in plants.*

| Group | K | Lambda | p K | p λ | Group | K | Lambda | p K | p λ |
| --- | --- | --- | --- | --- | --- | --- | --- | --- | --- |
| 24MC | 0.202 | 0.951 | 0.001 | 0.001 | ST(27:2)B | 0.135 | 0.963 | 0.013 | 0.001 |
| 24MD | 0.091 | 0.943 | 0.016 | 0.001 | ST(28:2)B | 0.363 | 0.818 | 0.003 | 0.001 |
| 24MP | 0.050 | 0.881 | 0.035 | 0.001 | ST(28:2)C | 0.210 | 0.762 | 0.003 | 0.001 |
| Agnosterol | 0.071 | 0.615 | 0.015 | 0.001 | ST(28:3)A | 0.059 | 0.667 | 0.026 | 0.002 |
| Anthelsterol | 0.091 | 0.585 | 0.007 | 0.001 | ST(28:3)B | 0.979 | 0.817 | 0.001 | 0.001 |
| Avenasterol | 0.047 | 0.579 | 0.009 | 0.001 | ST(29:0)C | 0.061 | 0.951 | 0.002 | 0.001 |
| β-Sitosterol | 0.101 | 0.965 | 0.001 | 0.001 | ST(29:1)A | 0.259 | 0.998 | 0.012 | 0.001 |
| Brassicasterol | 0.227 | 1.000 | 0.004 | 0.001 | ST(29:2)B | 0.112 | 1.000 | 0.010 | 0.001 |
| Campesterol | 0.226 | 0.977 | 0.001 | 0.001 | ST(30:2)E | 0.080 | 0.952 | 0.015 | 0.001 |
| Cycloartenol | 0.129 | 0.986 | 0.013 | 0.001 | ST(31:2)B | 0.190 | 0.997 | 0.029 | 0.001 |
| Desmosterol | 0.091 | 0.950 | 0.015 | 0.001 | Total sterol | 0.070 | 0.814 | 0.010 | 0.001 |
| Isofucosterol | 0.078 | 0.834 | 0.006 | 0.001 | 27 | 0.094 | 0.962 | 0.003 | 0.001 |
| Schottenol | 0.047 | 0.898 | 0.034 | 0.001 | 28 | 0.252 | 0.980 | 0.001 | 0.001 |
| Sitostanol | 0.060 | 0.848 | 0.045 | 0.001 | 29 | 0.100 | 0.949 | 0.001 | 0.001 |
| Spinasterol | 0.144 | 0.990 | 0.007 | 0.001 | 30 | 0.067 | 0.919 | 0.001 | 0.001 |
| Stigmasterol | 0.083 | 0.987 | 0.013 | 0.001 | CPR | 0.059 | 0.919 | 0.007 | 0.001 |
| ST(28:1)A | 0.104 | 0.797 | 0.023 | 0.001 | 0 | 0.060 | 0.848 | 0.036 | 0.001 |
| ST(27:1)A | 0.386 | 0.997 | 0.007 | 0.001 | 5 | 0.069 | 0.879 | 0.001 | 0.001 |
| ST(29:0)A | 0.306 | 1.000 | 0.018 | 0.001 | 7 | 0.042 | 0.761 | 0.015 | 0.001 |
| ST(29:2)D | 0.132 | 0.961 | 0.008 | 0.001 | NA | 0.033 | 0.724 | 0.047 | 0.001 |

***Table 6.*** *Summary of Indicator Species Analysis (ISA) results comparing the sterol profiles of all three Asteraceae subfamilies sampled (Asteroideae, Cichorioideae and Carduoideae – main text Figure 3). Indicator values (IV) and p-values are shown for all significant associations. All three subfamilies were associated with at least six un-named sterols demonstrating how poorly characterised the Asteraceae currently sterolome is. Carduoideae was associated with the most sterols and showed the smallest variation in sterolome between species. Data were averaged by species and subfamilies contained different number of species; Asteroideae (21), Carduoideae (14) and Cichorioideae (17).*

| Sterol | IV | P-value | Sterol | IV | P-value |
| --- | --- | --- | --- | --- | --- |
| Asteroideae | | | Carduoideae | | |
| Spinasterol | 0.783 | 0.005 | ST(27:2)F | 0.847 | 0.005 |
| ST(29:2)D | 0.738 | 0.005 | Lophenol | 0.759 | 0.005 |
| ST(29:3)B | 0.704 | 0.010 | ST(27:1)A | 0.757 | 0.005 |
| Sitostanol | 0.690 | 0.025 | ST(30:2)F | 0.739 | 0.005 |
| ST(28:0)E | 0.684 | 0.005 | Schottenol | 0.724 | 0.005 |
| ST(29:2)A | 0.683 | 0.010 | ST(29:3)A | 0.713 | 0.030 |
| ST(28:3)A | 0.663 | 0.045 | ST(29:0)C | 0.708 | 0.005 |
| Obtusifoliol | 0.660 | 0.025 | Cycloeucalenol | 0.705 | 0.010 |
| ST(28:3)B | 0.655 | 0.035 | ST(28:0)D | 0.682 | 0.005 |
| ST(28:0)A | 0.556 | 0.040 | ST(27:2)B | 0.674 | 0.010 |
| Cichorioideae | | | ST(31:2)A | 0.673 | 0.005 |
| ST(27:2)E | 0.912 | 0.005 | ST(28:2)B | 0.670 | 0.015 |
| Lathosterol | 0.860 | 0.005 | ST(30:2)E | 0.668 | 0.050 |
| Cycloartanol | 0.823 | 0.005 | ST(28:2)D | 0.663 | 0.030 |
| ST(30:2)B | 0.815 | 0.005 | ST(28:0)C | 0.663 | 0.025 |
| ST(27:2)G | 0.767 | 0.025 | ST(29:1)B | 0.646 | 0.020 |
| ST(29:1)C | 0.762 | 0.005 | ST(30:3) | 0.643 | 0.045 |
| ST(28:1)A | 0.755 | 0.005 | ST(30:2)A | 0.637 | 0.005 |
| ST(27:1)D | 0.740 | 0.045 |  |  |  |
| 24MC | 0.707 | 0.005 |  |  |  |
| Brassicasterol | 0.681 | 0.005 |  |  |  |
| Isofucosterol | 0.660 | 0.005 |  |  |  |

***Table 7.*** *Summary of Indicator Species Analysis (ISA) results on pollen sterol data comparing genera within families (Figure 4A). Out of 78 sterols tested, Asteraceae genera showed associations with 42, Fabaceae with 26 and Rosaceae with 38. Indicator values (IV) and p-values are shown for all. Data were analysed separately by family, each genus contained at least three species.*

| Sterol | IV | P-value | Sterol | IV | P-value | Sterol | IV | P-value |
| --- | --- | --- | --- | --- | --- | --- | --- | --- |
| Asteraceae | | | Fabaceae | | | Rosaceae | | |
| *Centaurea* | | | *Trifolium* | | | *Potentilla* | | |
| Cyclolaudenol | 0.804 | 0.005 | Schottenol | 0.856 | 0.005 | ST(27:2)D | 0.987 | 0.005 |
| Sitostanol | 0.801 | 0.005 | ST(27:2)C | 0.785 | 0.005 | ST(29:0)A | 0.952 | 0.005 |
| ST(28:0)E | 0.792 | 0.005 | 24MC | 0.754 | 0.005 | Schottenol | 0.940 | 0.005 |
| ST(28:2)B | 0.772 | 0.005 | Ergosterol | 0.670 | 0.015 | ST(28:3)A | 0.934 | 0.005 |
| ST(27:2)B | 0.736 | 0.005 | Spinasterol | 0.628 | 0.005 | 24MP | 0.928 | 0.005 |
| ST(29:2)A | 0.736 | 0.005 | *Ulex* | | | Brassicasterol | 0.875 | 0.020 |
| ST(30:3) | 0.734 | 0.005 | Sitostanol | 0.891 | 0.005 | ST(27:2)E | 0.849 | 0.015 |
| ST(27:1)E | 0.724 | 0.010 | ST(28:0)E | 0.888 | 0.005 | ST(27:1)C | 0.843 | 0.005 |
| ST(31:2)A | 0.701 | 0.005 | ST(30:2)E | 0.771 | 0.005 | Lathosterol | 0.814 | 0.005 |
| Ergosterol | 0.686 | 0.010 | ST(30:1) | 0.748 | 0.015 | Episterol | 0.789 | 0.025 |
| 24MCA | 0.674 | 0.015 | Stigmasterol | 0.748 | 0.020 | Spinasterol | 0.766 | 0.005 |
| 24MD | 0.655 | 0.040 | ST(29:1)D | 0.745 | 0.010 | ST(29:3)A | 0.765 | 0.005 |
| ST(30:1) | 0.650 | 0.030 | Cycloeucalenol | 0.742 | 0.005 | ST(29:2)C | 0.751 | 0.025 |
| ST(27:2)C | 0.636 | 0.035 | β-Sitosterol | 0.725 | 0.020 | ST(29:1)B | 0.744 | 0.005 |
| *Cirsium* | | | ST(30:2)B | 0.677 | 0.010 | Campesterol | 0.716 | 0.005 |
| ST(27:1)B | 0.916 | 0.005 | ST(29:0)C | 0.673 | 0.010 | ST(29:2)B | 0.704 | 0.045 |
| ST(27:2)F | 0.898 | 0.005 | ST(30:2)A | 0.664 | 0.005 | ST(29:1)A | 0.701 | 0.020 |
| ST(27:1)A | 0.810 | 0.005 | Campesterol | 0.652 | 0.035 | Cycloartenol | 0.692 | 0.020 |
| ST(29:0)B | 0.757 | 0.005 | ST(30:2)C | 0.641 | 0.010 | ST(29:0)C | 0.675 | 0.040 |
| ST(30:2)E | 0.713 | 0.010 | *Vicia* | | | *Prunus* | | |
| Cycloeucalenol | 0.700 | 0.005 | ST(29:1)C | 0.755 | 0.040 | ST(28:0)A | 0.840 | 0.010 |
| Spinasterol | 0.695 | 0.005 | ST(29:3)A | 0.737 | 0.005 | ST(31:2)A | 0.804 | 0.005 |
| ST(27:2)A | 0.694 | 0.005 | 24MCA | 0.729 | 0.015 | ST(28:2)B | 0.796 | 0.005 |
| Schottenol | 0.689 | 0.005 | Anthelsterol* | 0.716 | 0.005 | Cyclolaudenol | 0.791 | 0.005 |
| Lophenol | 0.686 | 0.010 | Cyclolaudenol | 0.696 | 0.005 | ST(28:3)B | 0.771 | 0.025 |
| 24MP | 0.677 | 0.015 | ST(31:2)A | 0.664 | 0.005 | Isofucosterol | 0.758 | 0.005 |
| ST(29:0)C | 0.677 | 0.005 | Cycloartenol | 0.620 | 0.030 | 24MC | 0.741 | 0.015 |
| ST(29:1)A | 0.664 | 0.045 | ST(28:0)B | 0.577 | 0.020 | β-Sitosterol | 0.731 | 0.005 |
| ST(30:2)B | 0.659 | 0.005 |  |  |  | ST(28:2)C | 0.708 | 0.025 |
| Avenasterol | 0.656 | 0.005 |  |  |  | ST(28:1)A | 0.707 | 0.045 |
| ST(30:2)C | 0.646 | 0.005 |  |  |  | Obtusifoliol | 0.691 | 0.040 |
| *Crepis* | | |  |  |  | *Rubus* | | |
| ST(28:3)B | 1.000 | 0.005 |  |  |  | Desmosterol | 0.992 | 0.005 |
| ST(27:2)E | 0.889 | 0.010 |  |  |  | ST(30:2)D | 0.918 | 0.005 |
| ST(28:1)A | 0.857 | 0.005 |  |  |  | Stigmasterol | 0.810 | 0.010 |
| ST(29:1)C | 0.846 | 0.010 |  |  |  | 24MCA | 0.804 | 0.010 |
| Cycloartanol | 0.833 | 0.005 |  |  |  | ST(30:2)E | 0.757 | 0.005 |
| 24MC | 0.791 | 0.005 |  |  |  | ST(27:2)A | 0.756 | 0.005 |
| ST(29:0)A | 0.777 | 0.010 |  |  |  | ST(27:1)E | 0.727 | 0.015 |
| Brassicasterol | 0.768 | 0.005 |  |  |  | ST(30:2)C | 0.666 | 0.005 |
| Obtusifoliol | 0.708 | 0.010 |  |  |  |  |  |  |
| Isofucosterol | 0.707 | 0.005 |  |  |  |  |  |  |
| Lathosterol | 0.706 | 0.005 |  |  |  |  |  |  |
| Cycloartenol | 0.674 | 0.020 |  |  |  |  |  |  |

***Table 8.*** *Summary of Indicator Species Analysis (ISA) results on pollen sterol data for the inter-species comparison shown in Figure 4B,C. Out of 78 sterols tested, 45 were significantly associated with a species group. Indicator values (IV) and p-values are shown for all 45. All species are associated with at least three sterols, indicating they have reliably distinct sterolomes as shown in Figure 4B,C.* Prunus spinosa *demonstrated the greatest variance in sterol profile and has the weakest sterol associations.*

| Sterol | IV | P-value | Sterol | IV | P-value |
| --- | --- | --- | --- | --- | --- |
| *Calluna vulgaris* | | | *Ranunculus ficaria* | | |
| Sitostanol | 0.805 | 0.005 | ST(29:2)B | 0.734 | 0.005 |
| ST(30:2)F | 0.638 | 0.040 | Isofucosterol | 0.627 | 0.005 |
| β-Sitosterol | 0.625 | 0.005 | ST(31:2)B | 0.580 | 0.045 |
| Stigmasterol | 0.592 | 0.005 | ST(30:2)A | 0.530 | 0.005 |
| ST(29:2)A | 0.555 | 0.005 | Obtusifoliol | 0.522 | 0.005 |
| ST(29:1)B | 0.545 | 0.005 | *Tripolium pannonicum* | | |
| *Hyacinthoides non-scripta* | | | ST(27:1)B | 0.991 | 0.005 |
| ST(29:3)A | 0.785 | 0.005 | ST(28:0)C | 0.758 | 0.010 |
| Schottenol | 0.773 | 0.005 | ST(27:1)E | 0.736 | 0.005 |
| ST(28:2)A | 0.703 | 0.005 | ST(30:1) | 0.687 | 0.005 |
| ST(28:1)A | 0.694 | 0.005 | ST(30:2)D | 0.668 | 0.010 |
| ST(29:2)D | 0.657 | 0.030 | ST(29:0)A | 0.643 | 0.005 |
| Cycloartanol | 0.607 | 0.005 | Spinasterol | 0.622 | 0.005 |
| *Knautia arvensis* | | | ST(27:2)B | 0.612 | 0.005 |
| 24MP | 0.644 | 0.005 | Avenasterol | 0.556 | 0.005 |
| ST(30:2)E | 0.599 | 0.010 | Cycloeucalenol | 0.555 | 0.010 |
| ST(27:1)C | 0.585 | 0.025 | ST(28:2)B | 0.552 | 0.005 |
| *Prunus spinosa* | | | ST(27:2)A | 0.549 | 0.005 |
| ST(28:3)B | 0.586 | 0.020 | ST(28:0)E | 0.534 | 0.005 |
| 24MC | 0.578 | 0.020 | ST(30:2)C | 0.474 | 0.005 |
| Cyclolaudenol | 0.498 | 0.005 |  |  |  |
| *Ranunculus acris* | | |  |  |  |
| Ergosterol | 0.826 | 0.005 |  |  |  |
| ST(28:3)A | 0.658 | 0.010 |  |  |  |
| ST(27:2)E | 0.627 | 0.005 |  |  |  |
| Cycloartenol | 0.576 | 0.005 |  |  |  |
| Campesterol | 0.572 | 0.005 |  |  |  |
| Lathosterol | 0.572 | 0.010 |  |  |  |
| 24MD | 0.518 | 0.010 |  |  |  |
| ST(29:1)A | 0.503 | 0.005 |  |  |  |

***Table 9.*** *Species with the highest and lowest total sterol content (mg/kg). Values were calculated as species means. Highlighted rows belong to the family Asteraceae and contain some of the highest and lowest values in the dataset. Brassicaceae species also showed consistently high total sterol production (*Lunaria annua, Cardamine pratensis, Brassica rapa*).*

| **Species** | **Total Sterol (mg/kg)** | **Species** | **Total Sterol (mg/kg)** |
| --- | --- | --- | --- |
| *Taraxacum sect. Taraxacum* | 75423.52 | *Malva sylvestris* | 729.98 |
| *Lapsana communis* | 63824.55 | *Centaurea cyanus* | 827.78 |
| *Lunaria annua* | 59551.78 | *Bellis perennis* | 831.20 |
| *Cardamine pratensis* | 52441.40 | *Carduus nutans* | 1072.10 |
| *Brassica rapa* | 38312.12 | *Pinus coulteri* | 1136.28 |

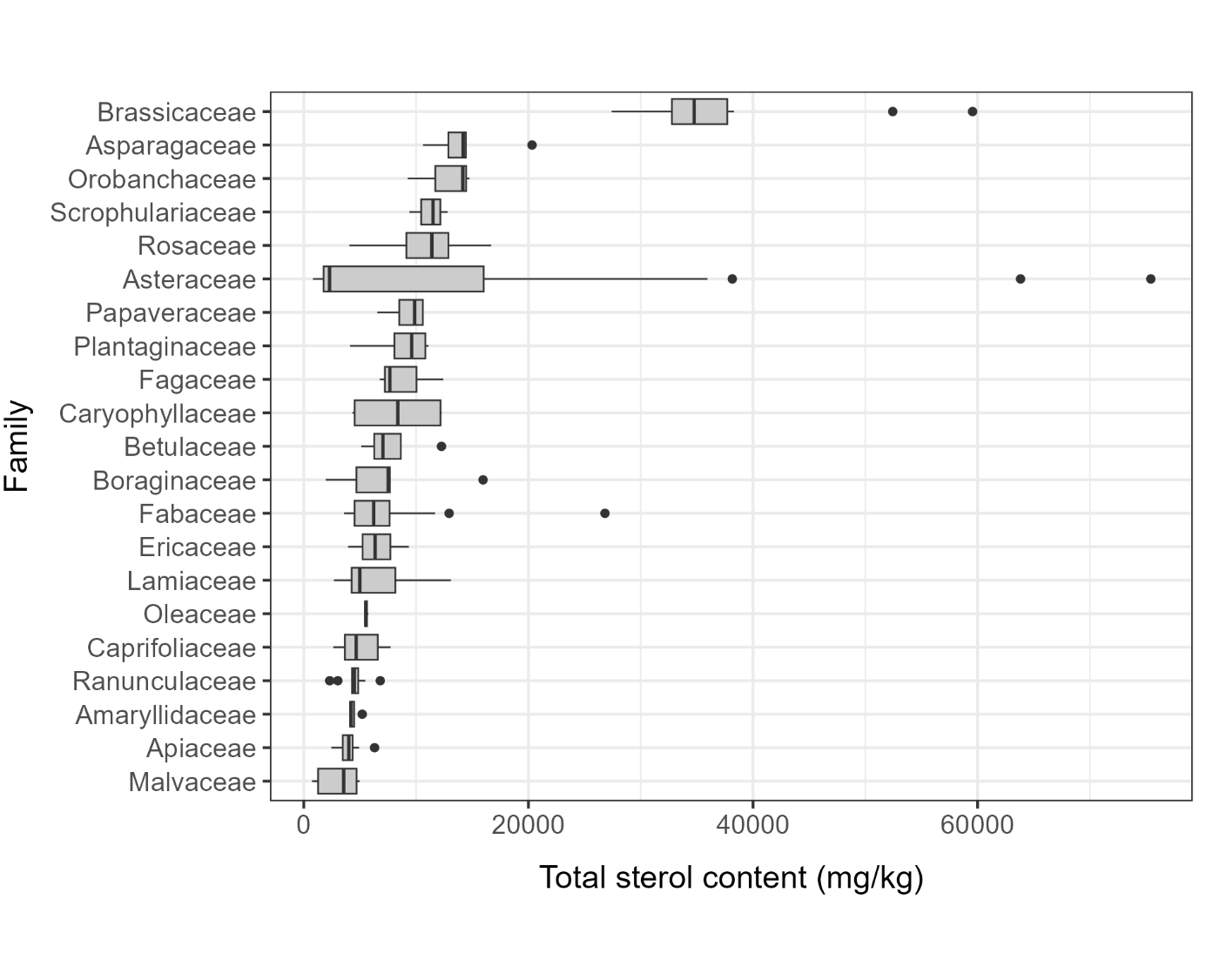

***Figure 4****. Total sterol production (mg/kg) ranged greatly within species samples. Graph shows the range in total sterol content across families containing at least three genera. Asteraceae is the most sampled family and also displays the highest range in sterol quantity. Brassicaceae shows a less variable high total sterol production. Boxes show first and third quartile, median and 1.5 x inter quartile range. Data is averaged by species, grouped by family and ranked by mean.*

| **Species** | **Total samples** | **Counties collected from** | **Mean (mg/kg)** | **SD (mg/kg)** | **Max (mg/kg)** | **Min (mg/kg)** |
| --- | --- | --- | --- | --- | --- | --- |
| *Acer pseudoplatanus* | 7 | 3 | 8610.56 | 1345.59 | 10928.20 | 6953.34 |
| *Aesculus hippocastanum* | 4 | 3 | 4604.56 | 547.78 | 5235.14 | 4002.78 |
| *Anemonoides nemorosa* | 4 | 3 | 2307.65 | 170.94 | 2524.54 | 2127.58 |
| *Arctium minus* | 4 | 3 | 2219.32 | 704.02 | 3265.20 | 1739.24 |
| *Betula pendula* | 5 | 3 | 5116.40 | 2029.07 | 6597.53 | 1589.77 |
| *Brassica napus* | 3 | 3 | 33451.56 | 3396.65 | 37013.05 | 30248.11 |
| *Calluna vulgaris* | 6 | 5 | 6348.89 | 1154.77 | 7264.60 | 4134.19 |
| *Centaurea scabiosa* | 3 | 3 | 1199.63 | 216.62 | 1449.55 | 1065.77 |
| *Cirsium vulgare* | 3 | 3 | 1709.68 | 211.23 | 1953.15 | 1575.38 |
| *Cornus sanguinea* | 3 | 3 | 3930.19 | 1241.60 | 5233.46 | 2761.16 |
| *Crepis capillaris* | 3 | 3 | 22428.86 | 3475.13 | 25239.07 | 18543.12 |
| *Cytisus scoparius* | 3 | 3 | 4110.61 | 972.25 | 5233.26 | 3544.54 |
| *Dipsacus fullonum* | 3 | 3 | 4412.75 | 449.28 | 4812.91 | 3926.74 |
| *Epilobium angustifolium* | 4 | 4 | 8474.81 | 1572.14 | 10182.08 | 6475.26 |
| *Erica tetralix* | 3 | 3 | 5238.58 | 1340.62 | 6386.28 | 3765.09 |
| *Filipendula ulmaria* | 4 | 4 | 8687.12 | 2123.70 | 11335.50 | 6174.78 |
| *Hyacinthoides non-scripta* | 9 | 4 | 14194.95 | 2261.07 | 16081.94 | 10057.01 |
| *Hyacinthoides x massartiana* | 3 | 3 | 12877.68 | 202.28 | 13110.38 | 12743.78 |
| *Hypochaeris radicata* | 4 | 3 | 6191.68 | 595.63 | 6762.11 | 5397.87 |
| *Impatiens glandulifera* | 3 | 3 | 2591.73 | 303.33 | 2939.30 | 2380.48 |
| *Jacobaea erucifolia* | 4 | 4 | 1870.75 | 460.03 | 2323.47 | 1283.10 |
| *Jacobaea vulgaris* | 4 | 4 | 1895.42 | 325.24 | 2235.33 | 1568.93 |
| *Knautia arvensis* | 6 | 4 | 3593.10 | 298.44 | 4050.03 | 3207.38 |
| *Leucanthemum vulgare* | 6 | 3 | 2171.35 | 207.51 | 2517.86 | 1977.68 |
| *Malus domestica* | 4 | 4 | 16512.81 | 3748.61 | 19858.41 | 11137.43 |
| *Malva moschata* | 5 | 3 | 1273.19 | 293.16 | 1523.88 | 783.99 |
| *Mercurialis perennis* | 5 | 3 | 9731.80 | 1206.43 | 11267.92 | 7884.40 |
| *Odontites vernus* | 3 | 3 | 14163.90 | 2713.54 | 16473.21 | 11175.23 |
| *Plantago coronopus* | 3 | 3 | 8646.77 | 1684.47 | 10471.89 | 7151.85 |
| *Plantago lanceolata* | 7 | 3 | 9205.73 | 1645.18 | 11387.39 | 6920.12 |
| *Primula vulgaris* | 8 | 3 | 7611.53 | 1654.30 | 10863.40 | 5842.85 |
| *Prunus avium* | 5 | 3 | 14347.66 | 5079.11 | 20662.50 | 9089.12 |
| *Prunus spinosa* | 5 | 4 | 12668.48 | 2747.69 | 14883.98 | 8061.50 |
| *Pulicaria dysenterica* | 4 | 4 | 2985.72 | 1141.45 | 4642.91 | 2208.35 |
| *Ranunculus acris* | 7 | 4 | 4619.54 | 659.70 | 5517.67 | 3768.29 |
| *Ranunculus bulbosus* | 6 | 3 | 4846.93 | 910.86 | 6033.13 | 4058.68 |
| *Ranunculus ficaria* | 8 | 4 | 5482.62 | 628.63 | 6584.11 | 4669.94 |
| *Ranunculus repens* | 4 | 3 | 4308.21 | 661.07 | 4985.55 | 3410.64 |
| *Reseda lutea* | 3 | 3 | 9977.46 | 2052.98 | 11566.78 | 7659.56 |
| *Rosa canina* | 4 | 3 | 6078.97 | 1957.45 | 8818.58 | 4485.81 |
| *Sanguisorba minor* | 4 | 3 | 12942.59 | 1427.02 | 14137.58 | 11224.99 |
| *Scorzoneroides autumnalis* | 3 | 3 | 15686.67 | 6467.89 | 23147.11 | 11656.49 |
| *Silene dioica* | 6 | 3 | 12283.42 | 804.68 | 13402.14 | 11328.09 |
| *Sonchus arvensis* | 3 | 3 | 38158.33 | 4914.28 | 41611.09 | 32532.08 |
| *Succisa pratensis* | 4 | 3 | 2616.96 | 208.29 | 2849.00 | 2434.76 |
| *Tripolium pannonicum* | 5 | 4 | 1308.60 | 618.60 | 1716.58 | 255.50 |
| *Ulex europaeus* | 5 | 3 | 7592.45 | 1999.13 | 9858.11 | 4941.57 |
| *Veronica chamaedrys* | 3 | 3 | 10931.14 | 1507.20 | 12666.23 | 9946.36 |

***Table 10.*** *Mean, standard deviation, maximum and minimum total sterol production (mg/kg) of species collected from at least three counties. Data shows large variation in total sterol production between species. Total sterol production also varied within species despite a largely consistent sterol profile. This intra-species variation was not found to be significantly influenced by collection county.*

| **Species** | **County collected** | **Month collected** | **Multi-site sample** | **Samples** |
| --- | --- | --- | --- | --- |
| *Acer campestre* | Oxfordshire | Apr-22 | No | 1 |
| *Acer platanoides* | Oxfordshire | Apr-21 | No | 1 |
| *Acer platanoides* | Sussex | Mar-22 | No | 1 |
| *Acer pseudoplatanus* | County Durham | Apr-22 | No | 1 |
| *Acer pseudoplatanus* | County Durham | May-22 | No | 2 |
| *Acer pseudoplatanus* | Greater London | Apr-22 | No | 2 |
| *Acer pseudoplatanus* | Oxfordshire | May-21 | No | 2 |
| *Acer sp.* | Oxfordshire | Apr-21 | No | 1 |
| *Achillea millefolium* | Oxfordshire | Aug-21 | Yes | 1 |
| *Achillea millefolium* | Oxfordshire | Jul-22 | Yes | 1 |
| *Achillea ptarmica* | Greater London | Jun-22 | No | 1 |
| *Aegopodium podagraria* | Greater London | May-22 | No | 1 |
| *Aegopodium podagraria* | Oxfordshire | Jun-22 | No | 1 |
| *Aegopodium podagraria* | Oxfordshire | May-22 | No | 1 |
| *Aesculus hippocastanum* | County Durham | May-22 | No | 1 |
| *Aesculus hippocastanum* | North Yorkshire | May-22 | No | 1 |
| *Aesculus hippocastanum* | Oxfordshire | May-21 | No | 2 |
| *Agrimonia eupatoria* | Oxfordshire & Sussex | Jul & Aug - 2021 (O), Jul - 2022 (S) | Yes | 1 |
| *Ajuga reptans* | Oxfordshire | May-21 | No | 1 |
| *Alcea rosea* | Oxfordshire | Aug-21 | No | 2 |
| *Alliaria petiolata* | Oxfordshire | Apr-22 | No | 1 |
| *Alliaria petiolata* | Oxfordshire | May-21 | No | 1 |
| *Allium triquetrum* | Oxfordshire | May-21 | No | 1 |
| *Allium ursinum* | North Yorkshire | May-22 | No | 1 |
| *Allium ursinum* | Oxfordshire | Apr-21 | No | 1 |
| *Allium ursinum* | Oxfordshire | May-21 | No | 1 |
| *Alnus glutinosa* | Greater London | Feb-22 | No | 3 |
| *Alnus glutinosa* | Oxfordshire | Feb-22 | No | 1 |
| *Alnus glutinosa* | Oxfordshire | Mar-21 | No | 5 |
| *Alnus glutinosa* | Oxfordshire | Mar-22 | No | 1 |
| *Anemonoides nemorosa* | Greater London | Mar-22 | No | 1 |
| *Anemonoides nemorosa* | Oxfordshire | Apr-22 | No | 1 |
| *Anemonoides nemorosa* | Oxfordshire | Mar-22 | Yes | 1 |
| *Anemonoides nemorosa* | Sussex | Mar-22 | No | 1 |
| *Anemonoides spp.* | West Yorkshire | Apr-21 | No | 1 |
| *Angelica sylvestris* | Oxfordshire | Aug-21 | No | 1 |
| *Anthriscus sylvestris* | Oxfordshire | Apr-22 | No | 1 |
| *Anthyllis vulneraria* | County Durham | Jun-22 | No | 1 |
| *Anthyllis vulneraria* | Sussex | Jul-22 | No | 1 |
| *Arctium agg.* | North Yorkshire | Jul-22 | No | 1 |
| *Arctium lappa* | Oxfordshire | Aug-21 | No | 2 |
| *Arctium lappa* | Oxfordshire | Jul-22 | No | 1 |
| *Arctium minus* | County Durham | Jul-22 | No | 1 |
| *Arctium minus* | Oxfordshire | Aug-21 | No | 1 |
| *Arctium minus* | Oxfordshire | Jul-22 | No | 1 |
| *Arctium minus* | Sussex | Jul-22 | No | 1 |
| *Argentina anserina* | Oxfordshire | May-21 | No | 1 |
| *Aria edulis* | Oxfordshire | May-21 | No | 1 |
| *Armeria maritima* | Dorset | Jul-22 | No | 1 |
| *Ballota nigra* | Oxfordshire | Jul-21 | No | 1 |
| *Ballota nigra* | Oxfordshire | Sep-21 | No | 1 |
| *Barbarea vulgaris* | Oxfordshire | May-21 | No | 1 |
| *Bellis perennis* | Oxfordshire | Feb-21 | No | 1 |
| *Bellis perennis* | Oxfordshire | Mar-21 | No | 2 |
| *Betula pendula* | County Durham | Apr-22 | No | 2 |
| *Betula pendula* | North Yorkshire | May-22 | No | 1 |
| *Betula pendula* | Oxfordshire | Apr-21 | No | 2 |
| *Borago officinalis* | Oxfordshire | Aug-21 | No | 3 |
| *Borago officinalis* | Oxfordshire | Sep-22 | No | 2 |
| *Brassica napus* | North Yorkshire | Apr-22 | No | 1 |
| *Brassica napus* | Oxfordshire | Apr-22 | No | 1 |
| *Brassica napus* | West Yorkshire | Jul-22 | No | 1 |
| *Brassica oleracea* | Oxfordshire | May-21 | No | 1 |
| *Brassica rapa* | Oxfordshire | May-22 | No | 1 |
| *Brassica sp.* | Oxfordshire | Jun-21 | No | 1 |
| *Brassica sp.* | Oxfordshire | May-21 | No | 1 |
| *Bryonia cretica* | Oxfordshire | Jun-21 | No | 1 |
| *Bryonia cretica subsp. Dioica* | Greater London | Jul-22 | No | 1 |
| *Buddleja davidii* | Oxfordshire | Aug-21 | No | 1 |
| *Buddleja davidii* | Oxfordshire | Sep-21 | No | 1 |
| *Calluna vulgaris* | Berkshire | Aug-22 | No | 1 |
| *Calluna vulgaris* | Dorset | Jul-22 | No | 1 |
| *Calluna vulgaris* | North Yorkshire | Aug-22 | No | 1 |
| *Calluna vulgaris* | North Yorkshire | Jul-22 | No | 1 |
| *Calluna vulgaris* | Oxfordshire | Sep-21 | No | 1 |
| *Calluna vulgaris* | Shropshire | Aug-21 | No | 1 |
| *Caltha palustris* | Oxfordshire | Apr-21 | No | 1 |
| *Calystegia sepium* | Greater London | Jun-22 | No | 1 |
| *Calystegia sepium* | North Yorkshire | Jul-22 | No | 1 |
| *Calystegia sepium/silvatica* | Oxfordshire | Aug-21 | No | 1 |
| *Calystegia sepium/silvatica* | Oxfordshire | Sep-21 | No | 1 |
| *Calystegia silvatica* | Oxfordshire | Sep-21 | No | 2 |
| *Calystegia sp.* | West Yorkshire | Jul-22 | No | 1 |
| *Campanula latifolia* | North Yorkshire | Jul-22 | No | 1 |
| *Campanula persicifolia* | Oxfordshire | Jun-21 | No | 1 |
| *Campanula rotundifolia* | Greater London | Jul-22 | No | 1 |
| *Campanula rotundifolia* | North Yorkshire | Jul-22 | No | 2 |
| *Campanula trachelium* | Oxfordshire | Aug-21 | No | 1 |
| *Cardamine pratensis* | Greater London | Mar-22 | No | 1 |
| *Cardamine pratensis* | Oxfordshire | Apr-21 | No | 1 |
| *Carduus crispus* | Oxfordshire | Jul-21 | No | 1 |
| *Carduus nutans* | Oxfordshire | Jun-21 | No | 2 |
| *Carpinus betulus* | Greater London | Mar-22 | No | 1 |
| *Carpinus betulus* | Oxfordshire | Apr-21 | No | 4 |
| *Carpinus betulus* | Oxfordshire | Mar-21 | No | 1 |
| *Castanea sativa* | Greater London | Jul-22 | No | 1 |
| *Centaurea cyanus* | Dorset | Jul-21 | No | 1 |
| *Centaurea montana* | Oxfordshire | Jun-21 | No | 1 |
| *Centaurea nigra* | County Durham | Jul-22 | No | 2 |
| *Centaurea nigra* | Oxfordshire | Jul-21 | No | 2 |
| *Centaurea nigra* | Oxfordshire | Jun-21 | No | 2 |
| *Centaurea scabiosa* | North Yorkshire | Jul-22 | No | 1 |
| *Centaurea scabiosa* | Oxfordshire | Jul-21 | No | 1 |
| *Centaurea scabiosa* | Wiltshire | Jul-21 | No | 1 |
| *Chelidonium majus* | Oxfordshire | May-21 | No | 1 |
| *Cichorium intybus* | Oxfordshire | Jul-21 | No | 1 |
| *Cichorium intybus* | Oxfordshire | Jul-21 | Yes | 1 |
| *Circaea lutetiana* | Oxfordshire | Jul - 2022 & Aug - 2022 | Yes | 1 |
| *Circaea lutetiana* | Oxfordshire & Greater London | Jul - 2021 (O), Jun - 2022 (GL) | Yes | 1 |
| *Cirsium acaule* | Sussex | Jul-22 | No | 2 |
| *Cirsium arvense* | County Durham | Jul-22 | No | 1 |
| *Cirsium arvense* | Oxfordshire | Jul-22 | No | 1 |
| *Cirsium arvense* | Oxfordshire | Jun-21 | No | 1 |
| *Cirsium eriophorum* | Oxfordshire | Aug-21 | No | 1 |
| *Cirsium eriophorum* | Oxfordshire | Aug-22 | No | 1 |
| *Cirsium eriophorum* | Wiltshire | Jul-21 | No | 1 |
| *Cirsium palustre* | North Yorkshire | Aug-22 | No | 1 |
| *Cirsium palustre* | North Yorkshire | Jul-22 | No | 1 |
| *Cirsium palustre* | Oxfordshire | Jul-21 | No | 1 |
| *Cirsium palustre* | Oxfordshire | Jul-22 | No | 1 |
| *Cirsium vulgare* | County Durham | Jul-22 | No | 1 |
| *Cirsium vulgare* | Greater London | May-22 | No | 1 |
| *Cirsium vulgare* | Oxfordshire | Jul-21 | No | 1 |
| *Clematis vitalba* | Essex | Jun-22 | No | 1 |
| *Clematis vitalba* | Oxfordshire | Jul-21 | No | 1 |
| *Clinopodium vulgare* | Oxfordshire | Aug-21 | No | 1 |
| *Convolvulus arvensis* | Greater London | Jun-22 | No | 1 |
| *Convolvulus arvensis* | North Yorkshire | Jul-22 | No | 1 |
| *Cornus sanguinea* | Greater London | May-22 | No | 1 |
| *Cornus sanguinea* | North Yorkshire | Jun-22 | No | 1 |
| *Cornus sanguinea* | Oxfordshire | May-22 | No | 1 |
| *Coronilla varia* | Oxfordshire | Sep-21 | No | 1 |
| *Corydalis solida* | Oxfordshire | Mar-21 | No | 2 |
| *Corylus avellana* | Greater London | Feb-22 | No | 1 |
| *Corylus avellana* | Greater London | Jan-22 | No | 4 |
| *Corylus avellana* | Oxfordshire | Feb-22 | No | 3 |
| *Corylus avellana* | Oxfordshire | Mar-22 | No | 1 |
| *Crataegus monogyna* | Greater London | May-22 | No | 1 |
| *Crataegus monogyna* | Oxfordshire | May-21 | No | 2 |
| *Crepis biennis* | Oxfordshire | May-21 | No | 1 |
| *Crepis capillaris* | County Durham | Jun-22 | No | 1 |
| *Crepis capillaris* | Oxfordshire | Jun-22 | No | 1 |
| *Crepis capillaris* | Sussex | Jun-22 | No | 1 |
| *Crepis vesicaria* | North Yorkshire | May-22 | No | 1 |
| *Crepis vesicaria* | Oxfordshire | Jun-21 | No | 1 |
| *Crocus sp.* | Oxfordshire | Mar-21 | No | 1 |
| *Crocus vernus* | Oxfordshire | Feb-21 | No | 2 |
| *Crocus vernus* | Oxfordshire | Mar-22 | No | 2 |
| *Crocus x luteus* | Oxfordshire | Feb-21 | No | 1 |
| *Cytisus scoparius* | County Durham | Apr-22 | No | 1 |
| *Cytisus scoparius* | North Yorkshire | May-22 | No | 1 |
| *Cytisus scoparius* | Tyne and Wear | Apr-22 | No | 1 |
| *Daucus carota* | Oxfordshire & North Yorkshire | Jul - 2021(O) , Jul - 2022 (NY) | Yes | 1 |
| *Digitalis purpurea* | County Durham | Jun-22 | No | 1 |
| *Digitalis purpurea* | Oxfordshire | Jul-21 | No | 1 |
| *Dioscorea communis* | Oxfordshire | Jun-22 | No | 1 |
| *Dipsacus fullonum* | County Durham | Jul-22 | No | 1 |
| *Dipsacus fullonum* | County Durham/ North Yorkshire | Aug-22 | No | 1 |
| *Dipsacus fullonum* | Oxfordshire | Sep-21 | No | 1 |
| *Dipsacus laciniatus* | Oxfordshire | Aug-21 | No | 1 |
| *Echium vulgare* | County Durham | Jun-22 | No | 1 |
| *Echium vulgare* | Oxfordshire | Jul-21 | No | 1 |
| *Echium vulgare* | Oxfordshire | Jun-21 | No | 1 |
| *Epilobium angustifolium* | County Durham | Jul-22 | No | 1 |
| *Epilobium angustifolium* | Greater London | Jul-22 | No | 1 |
| *Epilobium angustifolium* | Oxfordshire | Aug-21 | No | 1 |
| *Epilobium angustifolium* | Wiltshire | Jul-21 | No | 1 |
| *Epilobium hirsutum* | Greater London | Aug-22 | No | 1 |
| *Epilobium hirsutum* | West Yorkshire | Jul-22 | No | 1 |
| *Epilobium montanum* | Greater London | May-22 | No | 1 |
| *Epilobium parviflorum* | West Yorkshire & Sussex | Jun - 2022 (S), Jul - 2022 (WY) | Yes | 1 |
| *Erica cinerea* | Berkshire | Aug-22 | No | 1 |
| *Erica cinerea* | Dorset | Jul-22 | No | 1 |
| *Erica tetralix* | Berkshire | Aug-22 | No | 1 |
| *Erica tetralix* | Dorset | Jul-22 | No | 1 |
| *Erica tetralix* | North Yorkshire | Aug-22 | No | 1 |
| *Euonymus europaeus* | Oxfordshire | Apr & May - 2022 | No | 1 |
| *Eupatorium cannabinum* | Oxfordshire | Aug-21 | No | 2 |
| *Euphrasia officinalis* | County Durham | Aug-22 | No | 1 |
| *Fagus sylvatica* | Oxfordshire | Apr-21 | No | 2 |
| *Fallopia baldschuanica* | Oxfordshire | Jul-22 | No | 1 |
| *Filipendula ulmaria* | Greater London | Jul-22 | No | 1 |
| *Filipendula ulmaria* | Oxfordshire | Jul-21 | No | 1 |
| *Filipendula ulmaria* | Sussex | Jun-22 | No | 1 |
| *Filipendula ulmaria* | Tyne and Wear | Jun-22 | No | 1 |
| *Forsythia spp.* | West Yorkshire | Apr-21 | No | 1 |
| *Fraxinus excelsior* | Greater London | Apr-22 | No | 1 |
| *Fraxinus excelsior* | Oxfordshire | Apr-22 | No | 1 |
| *Fritillaria meleagris* | Oxfordshire | Apr-21 | No | 2 |
| *Galanthus nivalis* | Oxfordshire | Feb-21 | No | 2 |
| *Galanthus nivalis* | Oxfordshire | Feb-22 | No | 1 |
| *Galanthus nivalis* | Oxfordshire | Mar-21 | No | 2 |
| *Galanthus nivalis* | Oxfordshire | Mar-22 | No | 1 |
| *Galeopsis tetrahit* | Sussex | Aug-22 | No | 1 |
| *Galium verum* | Oxfordshire | Aug-22 | No | 1 |
| *Garrya elliptica* | Greater London | Jan-22 | No | 1 |
| *Geranium macrorrhizum* | Oxfordshire | May-21 | No | 1 |
| *Geranium pratense* | Oxfordshire | Jun-21 | No | 1 |
| *Geranium pyrenaicum* | Oxfordshire | May-21 | Yes | 1 |
| *Geranium robertianum* | Greater London | Apr-22 | No | 1 |
| *Geranium robertianum* | Oxfordshire | May-22 | No | 1 |
| *Geum urbanum* | County Durham & Greater London | May-22 | Yes | 1 |
| *Geum urbanum* | Oxfordshire | May-22 | No | 1 |
| *Glebionis segetum* | Dorset | Jul-21 | No | 1 |
| *Glechoma hederacea* | County Durham | Apr-22 | No | 1 |
| *Glechoma hederacea* | Oxfordshire | Apr-21 | No | 1 |
| *Glechoma hederacea* | Oxfordshire | May-21 | No | 1 |
| *Gunnera tinctoria var. tinctoria* | Greater London | NA | No | 1 |
| *Hedera helix* | Wiltshire, Oxfordshire & Greater London | Sep - 2021 (W & O), Sep - 2022 (GL) | Yes | 1 |
| *Helianthemum nummularium* | Oxfordshire | Jun-21 | Yes | 1 |
| *Helianthemum nummularium* | Oxfordshire | Sep-21 | No | 1 |
| *Helianthus annuus* | NA | Sep-21 | No | 4 |
| *Helleborus argutifolius* | Oxfordshire | Apr-21 | No | 1 |
| *Helminthotheca echioides* | Oxfordshire | Jul-21 | No | 1 |
| *Helminthotheca echioides* | Oxfordshire | Sep-22 | No | 1 |
| *Heracleum sphondylium* | Greater London | Jun-22 | No | 1 |
| *Heracleum sphondylium* | Oxfordshire | Jun-21 | No | 1 |
| *Hirschfeldia incana* | North Yorkshire | Jun-22 | No | 1 |
| *Hyacinthoides non-scripta* | County Durham | Apr-22 | No | 1 |
| *Hyacinthoides non-scripta* | Greater London | Apr-22 | No | 1 |
| *Hyacinthoides non-scripta* | North Yorkshire | Apr-22 | No | 1 |
| *Hyacinthoides non-scripta* | North Yorkshire | May-22 | No | 1 |
| *Hyacinthoides non-scripta* | Oxfordshire | Apr-21 | No | 2 |
| *Hyacinthoides non-scripta* | Oxfordshire | Apr-22 | No | 1 |
| *Hyacinthoides non-scripta* | Oxfordshire | May-21 | No | 2 |
| *Hyacinthoides x massartiana* | County Durham | Apr-22 | No | 1 |
| *Hyacinthoides x massartiana* | North Yorkshire | May-22 | No | 1 |
| *Hyacinthoides x massartiana* | Oxfordshire | Apr-21 | No | 1 |
| *Hypericum hirsutum* | Oxfordshire | Jul-21 | Yes | 1 |
| *Hypericum perforatum* | North Yorkshire | Aug & Sep - 2022 | No | 1 |
| *Hypericum perforatum* | Oxfordshire | Jul-21 | No | 1 |
| *Hypericum pulchrum* | Sussex | Jun-22 | No | 1 |
| *Hypochaeris radicata* | North Yorkshire | Jun-22 | No | 1 |
| *Hypochaeris radicata* | Oxfordshire | Jun-21 | No | 1 |
| *Hypochaeris radicata* | Oxfordshire | Jun-22 | No | 1 |
| *Hypochaeris radicata* | Oxfordshire & Greater London | Jun - 2021 (O), Jul - 2022 (GL) | Yes | 1 |
| *Ilex aquifolium* | Greater London | Apr-22 | No | 1 |
| *Ilex aquifolium* | Oxfordshire | May-22 | No | 1 |
| *Impatiens glandulifera* | County Durham | Sep-22 | No | 1 |
| *Impatiens glandulifera* | Greater London | Aug-22 | No | 1 |
| *Impatiens glandulifera* | Oxfordshire | Jul-21 | No | 1 |
| *Iris foetidissima* | Oxfordshire | Jun-22 | No | 1 |
| *Iris pseudacorus* | Greater London | May-22 | No | 2 |
| *Iris pseudacorus* | Oxfordshire | May-22 | No | 1 |
| *Jacobaea erucifolia* | County Durham | Aug-22 | No | 1 |
| *Jacobaea erucifolia* | Greater London | Jul-22 | No | 1 |
| *Jacobaea erucifolia* | Oxfordshire | Aug-22 | No | 1 |
| *Jacobaea erucifolia* | Wiltshire | Sep-21 | No | 1 |
| *Jacobaea vulgaris* | Greater London | Jun-22 | No | 1 |
| *Jacobaea vulgaris* | North Yorkshire | Jul-22 | No | 1 |
| *Jacobaea vulgaris* | Oxfordshire | Jun-22 | No | 1 |
| *Jacobaea vulgaris* | Sussex | Jul-22 | No | 1 |
| *Knautia arvensis* | County Durham | Jul-22 | No | 1 |
| *Knautia arvensis* | North Yorkshire | Jul-22 | No | 1 |
| *Knautia arvensis* | Oxfordshire | Jul-21 | No | 2 |
| *Knautia arvensis* | Oxfordshire | Jun-21 | No | 1 |
| *Knautia arvensis* | Wiltshire | Jul-21 | No | 1 |
| *Laburnum anagyroides* | Oxfordshire | May-21 | No | 1 |
| *Lactuca serriola* | Oxfordshire | Sep-21 | No | 1 |
| *Lamium album* | County Durham | Apr-22 | No | 1 |
| *Lamium album* | Oxfordshire | Apr-21 | No | 1 |
| *Lamium album* | Oxfordshire | Aug-21 | No | 2 |
| *Lamium galeobdolon* | Oxfordshire | Apr-21 | No | 1 |
| *Lamium galeobdolon subsp. Argentatum* | Oxfordshire | Apr-22 | No | 1 |
| *Lamium purpureum* | Greater London | Apr-22 | No | 1 |
| *Lapsana communis* | County Durham | Jun-22 | No | 1 |
| *Lapsana communis* | Oxfordshire | Jun-21 | Yes | 1 |
| *Lapsana communis* | Oxfordshire | Sep-21 | Yes | 1 |
| *Lathyrus latifolius* | Oxfordshire | Jul-21 | No | 2 |
| *Lathyrus pratensis* | Oxfordshire | Jul-21 | Yes | 1 |
| *Lathyrus pratensis* | Sussex | Jun-22 | No | 1 |
| *Leontodon hispidus* | Oxfordshire | Jun-22 | No | 1 |
| *Leontodon hispidus* | Oxfordshire | Sep-21 | No | 1 |
| *Leontodon hispidus* | Sussex | Jun-22 | No | 1 |
| *Leontodon saxatilis* | Greater London | Jun-22 | No | 1 |
| *Leontodon sp.* | Oxfordshire | Aug-21 | No | 1 |
| *Leucanthemum vulgare* | North Yorkshire | May-22 | No | 1 |
| *Leucanthemum vulgare* | Oxfordshire | Jul-21 | No | 1 |
| *Leucanthemum vulgare* | Oxfordshire | Jun-21 | No | 2 |
| *Leucanthemum vulgare* | Oxfordshire | May & Jun - 2021 | Yes | 1 |
| *Leucanthemum vulgare* | Sussex | Jun-22 | No | 1 |
| *Ligustrum sp.* | Oxfordshire | Jul-21 | No | 1 |
| *Ligustrum vulgare* | Oxfordshire | Jun-22 | No | 1 |
| *Lonicera caprifolium* | Oxfordshire | Jun-21 | No | 1 |
| *Lonicera periclymenum* | North Yorkshire | Aug-22 | No | 1 |
| *Lonicera periclymenum* | Oxfordshire | Jun-21 | No | 1 |
| *Lotus corniculatus* | North Yorkshire | Jun-22 | No | 1 |
| *Lotus corniculatus* | Oxfordshire | Aug-21 | No | 1 |
| *Lotus corniculatus* | Oxfordshire | Jul & Aug - 2021 | Yes | 1 |
| *Lotus corniculatus* | Oxfordshire | Jun-21 | No | 1 |
| *Lotus corniculatus* | Oxfordshire | Jun-21 | Yes | 1 |
| *Lotus corniculatus* | Oxfordshire | Sep-22 | No | 2 |
| *Lotus corniculatus subsp. Corniculatus* | North Yorkshire | May-22 | No | 1 |
| *Lotus pedunculatus* | North Yorkshire | Jul-22 | No | 1 |
| *Lotus pedunculatus* | Sussex | June & July - 2022 | Yes | 1 |
| *Lotus pedunculatus* | Sussex | Jun-22 | No | 1 |
| *Lunaria annua* | Oxfordshire | May-21 | No | 1 |
| *Lysimachia arvensis* | County Durham | Jun-22 | No | 1 |
| *Lysimachia nemorum* | Oxfordshire & Sussex | Jul - 2022 (O), Jun - 2022 (S) | Yes | 1 |
| *Lysimachia punctata* | Oxfordshire | Jul-21 | No | 1 |
| *Lysimachia vulgaris* | Oxfordshire | Aug-21 | No | 1 |
| *Lythrum salicaria* | Oxfordshire | Aug & Sep - 2021 | Yes | 1 |
| *Malus domestica* | County Durham | Apr-22 | No | 1 |
| *Malus domestica* | Greater London | Apr-22 | No | 1 |
| *Malus domestica* | North Yorkshire | Apr-22 | No | 1 |
| *Malus domestica* | Oxfordshire | May-21 | No | 1 |
| *Malus sp.* | Oxfordshire | Apr-22 | No | 3 |
| *Malva moschata* | Dorset | Jul-21 | No | 1 |
| *Malva moschata* | North Yorkshire | Jul-22 | No | 1 |
| *Malva moschata* | Oxfordshire | Jul-21 | No | 2 |
| *Malva moschata* | Oxfordshire | Jun-21 | No | 1 |
| *Malva sylvestris* | Greater London | Jun-22 | No | 1 |
| *Malva sylvestris* | Oxfordshire | Jun-21 | No | 4 |
| *Matricaria chamomilla* | County Durham | May-22 | No | 1 |
| *Matricaria chamomilla* | Oxfordshire | Aug-21 | No | 1 |
| *Medicago sativa* | Oxfordshire | Sep-21 | No | 1 |
| *Medicago sativa* | Oxfordshire | Sep-22 | No | 1 |
| *Melilotus officinalis* | Oxfordshire | Aug-21 | No | 1 |
| *Melilotus sp.* | Oxfordshire | Aug-21 | No | 2 |
| *Mentha aquatica* | County Durham | Aug-22 | No | 1 |
| *Mentha aquatica* | Oxfordshire | Aug-21 | No | 1 |
| *Mercurialis perennis* | Greater London | Apr-22 | No | 1 |
| *Mercurialis perennis* | North Yorkshire | Apr-22 | No | 2 |
| *Mercurialis perennis* | Oxfordshire | Apr-21 | No | 2 |
| *Muscari armeniacum* | Wiltshire | Apr-21 | No | 1 |
| *Narcissus pseudonarcissus* | Greater London | Feb-22 | No | 1 |
| *Narcissus sp.* | Oxfordshire | Apr-21 | No | 1 |
| *Narcissus sp.* | Oxfordshire | Feb-21 | No | 3 |
| *Narcissus sp.* | Oxfordshire | Mar-21 | No | 4 |
| *Narcissus sp.* | Oxfordshire | Mar-22 | No | 2 |
| *Narthecium ossifragum* | Northumberland | Jul-22 | No | 1 |
| *Nigella damascena* | Wiltshire | Sep-21 | No | 1 |
| *Odontites vernus* | Oxfordshire | Aug-21 | No | 1 |
| *Odontites vernus* | Wiltshire | Jul-21 | No | 1 |
| *Odontites vernus* | Wiltshire & Oxfordshire | Aug - 2021 (W & O) | Yes | 1 |
| *Oenanthe crocata* | Greater London | May-22 | No | 1 |
| *Oenanthe crocata* | Oxfordshire | May-22 | No | 1 |
| *Onobrychis viciifolia* | County Durham | Jul-22 | No | 1 |
| *Onobrychis viciifolia* | Wiltshire | Aug-21 | No | 1 |
| *Onobrychis viciifolia* | Wiltshire | Jul-21 | No | 1 |
| *Ononis spinosa subsp. Procurrens* | North Yorkshire | Jun-22 | No | 1 |
| *Ononis spinosa subsp. Procurrens* | Oxfordshire | Sep-21 | No | 1 |
| *Onopordum acanthium* | Oxfordshire | Jul-21 | No | 1 |
| *Origanum vulgare* | Oxfordshire | Aug-21 | No | 1 |
| *Oxalis acetosella* | North Yorkshire | May-22 | No | 1 |
| *Papaver cambricum* | Oxfordshire | Jun-21 | No | 1 |
| *Papaver cambricum* | Oxfordshire | May-21 | No | 1 |
| *Papaver rhoeas* | Greater London | Jun-22 | No | 1 |
| *Papaver rhoeas* | Oxfordshire | Jun-21 | No | 1 |
| *Passiflora caerulea* | Oxfordshire | Aug-21 | No | 1 |
| *Pastinaca sativa* | County Durham | Jul-22 | No | 1 |
| *Pastinaca sativa* | Oxfordshire | Jul-22 | No | 1 |
| *Petasites hybridus* | North Yorkshire | Apr-22 | No | 2 |
| *Petasites hybridus* | Oxfordshire | Apr-22 | No | 2 |
| *Petasites pyrenaicus* | Oxfordshire | Feb-22 | No | 1 |
| *Phacelia tanacetifolia* | Oxfordshire | Sep-21 | No | 1 |
| *Picris hieracioides* | Oxfordshire | Aug-21 | No | 1 |
| *Pilosella aurantiaca* | County Durham | Jun-22 | No | 1 |
| *Pilosella officinarum* | Oxfordshire | Jun-22 | No | 1 |
| *Pilosella officinarum* | Oxfordshire, County Durham & Greater London | Aug - 2021 (O), Jun - 2022 (CD), May - 2022 (GL) | Yes | 1 |
| *Pinus coulteri* | Greater London | 2022 | No | 1 |
| *Plantago coronopus* | Dorset | Jul-22 | No | 1 |
| *Plantago coronopus* | Greater London | May-22 | No | 1 |
| *Plantago coronopus* | North Yorkshire | Jun-22 | No | 1 |
| *Plantago lanceolata* | Greater London | May-22 | No | 1 |
| *Plantago lanceolata* | North Yorkshire | May-22 | No | 2 |
| *Plantago lanceolata* | Oxfordshire | Aug-21 | No | 1 |
| *Plantago lanceolata* | Oxfordshire | May-21 | No | 3 |
| *Plantago major* | Oxfordshire | Jul-21 | No | 1 |
| *Plantago maritima* | County Durham | Jun-22 | No | 1 |
| *Plantago maritima* | Dorset | Jul-22 | No | 1 |
| *Plantago media* | Oxfordshire | Jun & Jul - 2021 | Yes | 1 |
| *Plantago media* | Oxfordshire | Jun-21 | No | 1 |
| *Plantago media* | Oxfordshire | Jun - 2021, Aug - 2021 | Yes | 1 |
| *Populus tremula* | Oxfordshire | Mar-22 | Yes | 1 |
| *Potentilla erecta* | Berkshire | May-22 | No | 1 |
| *Potentilla reptans* | Oxfordshire | Jun-22 | No | 1 |
| *Potentilla sterilis* | Oxfordshire & North Yorkshire | Apr - 2022 (O), Mar - 2022 (NY) | Yes | 1 |
| *Primula veris* | Oxfordshire | Apr-21 | No | 2 |
| *Primula vulgaris* | County Durham & North Yorkshire | Apr - 2022 (CD & NY) | Yes | 1 |
| *Primula vulgaris* | Oxfordshire | Feb-21 | No | 1 |
| *Primula vulgaris* | Oxfordshire | Mar-21 | No | 4 |
| *Primula vulgaris* | Oxfordshire | Mar-22 | No | 1 |
| *Primula vulgaris* | Sussex | Mar-22 | No | 1 |
| *Prunella vulgaris* | Oxfordshire | Jun-22 | No | 1 |
| *Prunus avium* | Greater London | Mar-22 | No | 1 |
| *Prunus avium* | Oxfordshire | Apr-21 | No | 2 |
| *Prunus avium* | Oxfordshire | Mar-21 | No | 1 |
| *Prunus avium* | Sussex | Mar-22 | No | 1 |
| *Prunus cerasifera* | Oxfordshire | Mar-21 | No | 6 |
| *Prunus cerasifera* | Oxfordshire | Mar-22 | No | 1 |
| *Prunus spinosa* | Berkshire | Apr-21 | No | 1 |
| *Prunus spinosa* | County Durham | Apr-22 | Yes | 1 |
| *Prunus spinosa* | Oxfordshire | Apr-21 | No | 1 |
| *Prunus spinosa* | Oxfordshire | Mar-21 | Yes | 1 |
| *Prunus spinosa* | Sussex | Mar-22 | No | 1 |
| *Pulicaria dysenterica* | Oxfordshire | Aug-21 | No | 1 |
| *Pulicaria dysenterica* | Oxfordshire & North Yorkshire | Sep - 2021 (O), Sep - 2022 (NY) | Yes | 1 |
| *Pulicaria dysenterica* | Sussex | Aug-22 | No | 1 |
| *Pulicaria dysenterica* | Wiltshire | Sep-21 | No | 1 |
| *Pyrus sp.* | Oxfordshire | Apr-21 | No | 1 |
| *Quercus robur* | Greater London | Apr-22 | No | 1 |
| *Rabelera holostea* | North Yorkshire | Apr-22 | No | 1 |
| *Rabelera holostea* | North Yorkshire | May-22 | No | 1 |
| *Rabelera holostea* | Oxfordshire | Apr-22 | No | 1 |
| *Rabelera holostea* | Oxfordshire | May-21 | No | 3 |
| *Ranunculus acris* | County Durham | May-22 | No | 1 |
| *Ranunculus acris* | Greater London | May-22 | No | 2 |
| *Ranunculus acris* | North Yorkshire | Jun-22 | No | 1 |
| *Ranunculus acris* | North Yorkshire | May-22 | No | 1 |
| *Ranunculus acris* | Oxfordshire | Jun-21 | No | 1 |
| *Ranunculus acris* | Oxfordshire | May-21 | No | 1 |
| *Ranunculus bulbosus* | Greater London | Apr-22 | No | 1 |
| *Ranunculus bulbosus* | Greater London | NA | No | 1 |
| *Ranunculus bulbosus* | North Yorkshire | May-22 | No | 1 |
| *Ranunculus bulbosus* | Oxfordshire | Apr-21 | No | 2 |
| *Ranunculus bulbosus* | Oxfordshire | May-21 | No | 1 |
| *Ranunculus ficaria* | Greater London | Mar-22 | No | 1 |
| *Ranunculus ficaria* | North Yorkshire | Apr-22 | No | 1 |
| *Ranunculus ficaria* | North Yorkshire | Mar-22 | No | 1 |
| *Ranunculus ficaria* | Oxfordshire | Mar-21 | No | 4 |
| *Ranunculus ficaria* | Wiltshire | Mar-22 | No | 1 |
| *Ranunculus repens* | Greater London | May-22 | No | 1 |
| *Ranunculus repens* | North Yorkshire | Jun-22 | No | 1 |
| *Ranunculus repens* | Oxfordshire | Jul-21 | No | 1 |
| *Ranunculus repens* | Oxfordshire | May-21 | No | 1 |
| *Ranunculus sp.* | Oxfordshire | Mar-21 | No | 1 |
| *Raphanus raphanistrum* | Greater London | Jun-22 | No | 1 |
| *Raphanus raphanistrum subsp. Sativus* | Greater London | Sep-22 | No | 1 |
| *Reseda lutea* | County Durham | Jul-22 | No | 1 |
| *Reseda lutea* | Greater London | May-22 | No | 1 |
| *Reseda lutea* | North Yorkshire | Jun-22 | No | 1 |
| *Reseda luteola* | Greater London | May-22 | No | 1 |
| *Reseda luteola* | Tyne and Wear | May-22 | No | 1 |
| *Rhamphospermum arvense* | North Yorkshire | May-22 | No | 1 |
| *Rhamphospermum arvense* | Oxfordshire | Apr-22 | No | 1 |
| *Rhamphospermum arvense* | Oxfordshire | Jun-22 | No | 1 |
| *Rhinanthus minor* | North Yorkshire | Jun-22 | No | 1 |
| *Rhinanthus minor* | Oxfordshire | Jul-21 | No | 1 |
| *Rhinanthus minor* | Oxfordshire | Jun-21 | No | 2 |
| *Rhododendron ponticum* | Greater London | May-22 | No | 1 |
| *Rhododendron ponticum* | Oxfordshire | Jun-21 | No | 1 |
| *Rosa arvensis* | Sussex | Jun-22 | No | 2 |
| *Rosa canina* | Greater London | May-22 | No | 1 |
| *Rosa canina* | North Yorkshire | Jun-22 | No | 1 |
| *Rosa canina* | Oxfordshire | Jun-21 | Yes | 1 |
| *Rosa canina* | Oxfordshire | Jun-22 | No | 1 |
| *Rubus caesius* | Wiltshire | Sep-21 | No | 1 |
| *Rubus fruticosus* | Oxfordshire | Aug-21 | No | 1 |
| *Rubus fruticosus* | Oxfordshire | Jun-21 | No | 1 |
| *Rubus fruticosus* | Wiltshire | Jul-21 | No | 1 |
| *Rubus idaeus* | Oxfordshire | Jun-22 | No | 1 |
| *Rubus idaeus* | Sussex | Jun-22 | No | 1 |
| *Salix caprea* | Oxfordshire | Mar-21 | No | 1 |
| *Salix caprea* | Oxfordshire | Mar-22 | No | 1 |
| *Salix caprea â€œKilmarnockâ€* | Greater London | Mar-22 | No | 1 |
| *Salix caprea/cinerea* | Berkshire | Apr-21 | No | 1 |
| *Salix caprea/cinerea* | County Durham | Apr-22 | No | 3 |
| *Salix caprea/cinerea* | Oxfordshire | Mar-21 | No | 4 |
| *Salix caprea/cinerea* | Sussex | Mar-22 | No | 1 |
| *Salix fragilis/alba* | Greater London | Apr-22 | No | 1 |
| *Salix repens* | Greater London | Apr-22 | No | 1 |
| *Salix sp.* | County Durham | Apr-22 | No | 1 |
| *Salix sp.* | Oxfordshire | Apr-22 | No | 2 |
| *Salix sp.* | Oxfordshire | Mar-21 | No | 2 |
| *Salix x pendulina nothof. Tristis* | Oxfordshire | Apr-21 | No | 2 |
| *Sambucus nigra* | Oxfordshire | Jun-21 | No | 3 |
| *Sanguisorba minor* | County Durham | Jun-22 | No | 1 |
| *Sanguisorba minor* | Greater London | Jun-22 | No | 2 |
| *Sanguisorba minor* | Oxfordshire | Apr-22 | No | 1 |
| *Scabiosa columbaria* | Wiltshire | Jul-21 | No | 1 |
| *Scilla forbesii* | Oxfordshire | Mar-21 | No | 2 |
| *Scilla siberica* | Oxfordshire | Mar-21 | No | 1 |
| *Scorzoneroides autumnalis* | County Durham | Sep-22 | No | 1 |
| *Scorzoneroides autumnalis* | Greater London | Jul-22 | No | 1 |
| *Scorzoneroides autumnalis* | Oxfordshire | Aug-21 | Yes | 1 |
| *Scrophularia vernalis* | Oxfordshire | May-21 | No | 1 |
| *Senecio vulgaris* | Greater London & North Yorkshire | Jun - 2022 (NY), Aug - 2022 (GL) | Yes | 1 |
| *Senecio/Jacobea sp.* | Oxfordshire | Jul-21 | No | 2 |
| *Silene dioica* | County Durham | May-22 | No | 2 |
| *Silene dioica* | North Yorkshire | May-22 | No | 1 |
| *Silene dioica* | Oxfordshire | Jun-21 | No | 1 |
| *Silene dioica* | Oxfordshire | Jun, Jul & Aug - 2021 | Yes | 1 |
| *Silene dioica* | Oxfordshire | May-21 | No | 1 |
| *Silene latifolia* | Oxfordshire | Aug-21 | No | 1 |
| *Silene latifolia* | Oxfordshire | Jun-21 | No | 1 |
| *Solanum dulcamara* | Greater London | May-22 | No | 1 |
| *Solanum dulcamara* | Oxfordshire | Jun-22 | No | 1 |
| *Solanum lycopersicum* | NA | Jul-21 | No | 1 |
| *Solanum nigrum* | Greater London | Jun-22 | No | 1 |
| *Solanum tuberosum* | North Yorkshire | Jul-22 | No | 1 |
| *Sonchus arvensis* | County Durham | Aug-22 | No | 1 |
| *Sonchus arvensis* | North Yorkshire | Jun-22 | No | 1 |
| *Sonchus arvensis* | Sussex | Jul-22 | No | 1 |
| *Sonchus oleraceus* | Greater London | May-22 | No | 1 |
| *Sonchus oleraceus* | Oxfordshire, County Durham & North Yorkshire | Jun - 2022 (NY), Jul - 2022 (CD), Jul - 2022 (O) | Yes | 1 |
| *Sorbus aucuparia* | County Durham | May-22 | No | 1 |
| *Sorbus aucuparia* | Oxfordshire | May-21 | No | 3 |
| *Stachys sylvatica* | Oxfordshire | Jun-22 | No | 1 |
| *Stachys sylvatica* | Sussex | Jun-22 | No | 1 |
| *Stellaria graminea* | Oxfordshire | Jun-22 | No | 1 |
| *Succisa pratensis* | County Durham | Aug-22 | No | 1 |
| *Succisa pratensis* | Greater London | Sep-22 | No | 1 |
| *Succisa pratensis* | Oxfordshire | Aug-21 | No | 1 |
| *Succisa pratensis* | Oxfordshire | Sep-21 | No | 1 |
| *Symphyotrichum lanceolatum* | Oxfordshire | Sep-21 | No | 2 |
| *Symphyotrichum x salignum* | County Durham | Sep-22 | No | 1 |
| *Symphytum orientale* | Oxfordshire | Apr-21 | No | 1 |
| *Symphytum sp.* | Oxfordshire | Jun-21 | No | 1 |
| *Symphytum x uplandicum* | Greater London | May-22 | No | 1 |
| *Tanacetum vulgare* | County Durham | Jul-22 | No | 1 |
| *Tanacetum vulgare* | County Durham | Sep-22 | No | 1 |
| *Tanacetum vulgare* | North Yorkshire | Jul-22 | No | 1 |
| *Taraxacum sect. Taraxacum* | Greater London | Apr-22 | No | 1 |
| *Taraxacum sect. Taraxacum* | Oxfordshire | Apr-21 | No | 1 |
| *Teucrium scorodonia* | Sussex | Jun-22 | No | 1 |
| *Tilia platyphyllos* | Greater London | Jun-22 | No | 3 |
| *Tilia x europaea* | Oxfordshire | Jun-21 | No | 1 |
| *Torilis japonica* | North Yorkshire | Jul-22 | No | 1 |
| *Torminalis glaberrima* | Greater London | May-22 | No | 1 |
| *Trifolium hybridum* | Oxfordshire | Sep-22 | No | 1 |
| *Trifolium incarnatum* | Oxfordshire | Sep-22 | No | 3 |
| *Trifolium pratense* | Oxfordshire | Jul-21 | No | 4 |
| *Trifolium pratense* | Oxfordshire | Jul-21 | Yes | 1 |
| *Trifolium pratense* | Oxfordshire | Sep-21 | No | 1 |
| *Trifolium pratense* | Oxfordshire | Sep-22 | No | 3 |
| *Trifolium repens* | Oxfordshire | Aug-21 | No | 3 |
| *Trifolium repens* | Oxfordshire | Sep-22 | No | 1 |
| *Trifolium resupinatum* | Oxfordshire | Sep-22 | No | 2 |
| *Tripleurospermum inodorum* | Oxfordshire | Jul-21 | No | 1 |
| *Tripleurospermum inodorum* | Oxfordshire | Sep-22 | No | 1 |
| *Tripleurospermum maritimum* | Dorset | Jul-22 | No | 1 |
| *Tripolium pannonicum* | County Durham | Aug-22 | No | 1 |
| *Tripolium pannonicum* | County Durham/North Yorkshire | Aug-22 | No | 1 |
| *Tripolium pannonicum* | Essex | Sep-22 | No | 2 |
| *Tripolium pannonicum* | Kent | Sep-22 | No | 1 |
| *Tulipa sylvestris* | Oxfordshire | Apr-21 | No | 3 |
| *Tussilago farfara* | County Durham | Mar-22 | No | 1 |
| *Ulex europaeus* | Greater London | Mar-22 | No | 1 |
| *Ulex europaeus* | Oxfordshire | Mar-22 | No | 1 |
| *Ulex europaeus* | Oxfordshire | May-21 | No | 2 |
| *Ulex europaeus* | Sussex | Mar-22 | No | 1 |
| *Ulex gallii* | Dorset | Sep-22 | No | 1 |
| *Ulex minor* | Berkshire & Dorset | Aug - 2022 (B), Sep - 2022 (D) | Yes | 1 |
| *Ulex minor* | Hampshire | Aug-22 | No | 1 |
| *Vaccinium myrtillus* | Berkshire | May-22 | No | 1 |
| *Vaccinium myrtillus* | North Yorkshire | May-22 | No | 2 |
| *Valeriana rubra* | Oxfordshire | Jun-21 | No | 2 |
| *Verbascum thapsus* | Oxfordshire | Jul - 2021, Aug - 2021 | Yes | 1 |
| *Veronica chamaedrys* | Greater London | May-22 | No | 1 |
| *Veronica chamaedrys* | North Yorkshire | May-22 | No | 1 |
| *Veronica chamaedrys* | Oxfordshire | May-21 | No | 1 |
| *Veronica filiformis* | Oxfordshire | Apr-22 | No | 1 |
| *Veronica montana* | Oxfordshire | May-22 | No | 1 |
| *Veronica officinalis* | Oxfordshire | Jun-22 | No | 1 |
| *Veronica officinalis* | Sussex | Jun-22 | No | 1 |
| *Veronica persica* | Oxfordshire | Mar-21 | No | 1 |
| *Viburnum lantana* | Oxfordshire | Apr-21 | No | 1 |
| *Viburnum lantana* | Oxfordshire | May-21 | No | 1 |
| *Vicia cracca* | Oxfordshire | Jul - 2021, Aug - 2021, Jul - 2022, Aug - 2022 | Yes | 1 |
| *Vicia sativa* | Oxfordshire | May-22 | No | 1 |
| *Vicia sativa* | Oxfordshire, County Durham & North Yorkshire | May - 2021 (O), Apr - 2022 (O), Jun - 2022 (NY), May - 2022 (CD) | Yes | 1 |
| *Vicia sepium* | North Yorkshire | Apr-22 | No | 1 |
| *Vicia sepium* | Oxfordshire | May-22 | No | 2 |
| *Viola odorata* | Oxfordshire | Mar-21 | No | 1 |
| *Viola odorata* | Oxfordshire | Mar - 2021, Mar - 2022 | Yes | 1 |
| *Viola riviniana* | North Yorkshire | Apr-22 | No | 1 |
| *Viola riviniana* | Sussex | Apr-22 | No | 1 |

***Table 11****. Collection details for all pollen samples. Multi-site samples show where multiple under-weight samples were combined to reach minimum analysis weight of 10 mg. In these cases, collection month and year are given for each county which formed part of the composite sample. Samples were extracted by collected from 15 counties across February 2021 to September 2022.*

| **Sterol** | **Structure** | **m/z** | **Retention time (min)** |
| --- | --- | --- | --- |
| 24-methylenecholesterol (24MC) | (28:2) | 398.3549 | 5.9 |
| 24-methylenecycloartenol (24MCA) | (31:2) | 440.4018 | 7.5 |
| Anthelsterol | (29:3) | 410.3549 | 5.3 |
| Avenasterol | (29:2) | 412.3705 | 6.9 |
| β-sitosterol | (29:1) | 414.3862 | 9.7 |
| Brassicasterol | (28:2) | 398.3549 | 6.6 |
| Campesterol | (28:1) | 400.3705 | 8.2 |
| Cholesterol | (27:1) | 386.3549 | 7.0 |
| Cycloartenol | (30:2) | 426.3862 | 8.3 |
| Cycloeucalenol | (30:2) | 426.3862 | 7.7 |
| Cyclolaudenol | (31:2) | 440.4018 | 9.5 |
| Desmosterol | (27:2) | 384.3392 | 5.1 |
| Episterol | (28:2) | 398.3549 | 5.7 |
| Ergosterol | (28:3) | 396.3392 | 5.5 |
| Isofucosterol | (29:2) | 412.3705 | 7.5 |
| Schottenol | (29:1) | 414.3862 | 9.0 |
| Sitostanol | (29:0) | 416.4018 | 11.1 |
| Spinasterol | (29:2) | 412.3705 | 8.0 |
| Stigmasterol | (29:2) | 412.3705 | 8.5 |

***Table 12.*** *Retention time, mass to charge ratio (m/z) and structure description of sterols (carbon count (CC): double bond equivalents (DB)) used in Quality Assurance (QA) mixture.*

| **Sterol** | **Carbons** | **Double bond** |
| --- | --- | --- |
| Cholesterol | 27 | 5 |
| Lathosterol | 27 | 7 |
| Desmosterol | 27 | 5 |
| Campesterol | 28 | 5 |
| Lophenol | 28 | 7 |
| 24MC | 28 | 5 |
| 24MD | 28 | 5 |
| Brassicasterol | 28 | 5 |
| Episterol | 28 | 7 |
| Ergosterol | 28 | NA |
| Sitostanol | 29 | 0 |
| β-Sitosterol | 29 | 5 |
| Schottenol | 29 | 7 |
| 24MP | 29 | CPR |
| Avenasterol | 29 | 7 |
| Isofucosterol | 29 | 5 |
| Spinasterol | 29 | 7 |
| Stigmasterol | 29 | 5 |
| Cycloartanol | 30 | CPR |
| 24MCA | 30 | CPR |
| Cycloartenol | 30 | CPR |
| Cycloeucalenol | 30 | CPR |
| Obtusifoliol | 30 | 8 |
| Cyclolaudenol | 31 | CPR |

***Table 13.*** *Summary of carbon count double bond positions of sterols identified using reference materials. Full structure and formula for each sterol, except ergosterol, is shown in Figure 2. Ergosterol is classed as NA due to it containing double bonds at positions five and seven in the steroid nucleus B-ring. CPR = Cyclopropane ring.*
